# Transglutaminase 2 Deletion Enhances Astrocyte-to-Neuron Metabolic Support and Attenuates Subacute Pathology Following Repetitive Mild Traumatic Brain Injury

**DOI:** 10.64898/2026.07.16.738974

**Authors:** Thomas Delgado, Tanzil M. Arefin, Ian Pagan, Benjamin H. Weekley, Joel Rodwell-Bullock, Mohammad R. Chowdhury, Anthony B. Crum, Ian Maze, Paul S. Brookes, Julian P. Meeks, Gail V.W. Johnson

## Abstract

Mild traumatic brain injury (mTBI) is the most common form of central nervous system (CNS) injury and is often characterized by persistent neuroinflammation, metabolic dysregulation, and oxidative stress. Repetitive injuries compound these pathologies and lead to multifocal axonal injuries and long-term functional deficits. Despite the prevalence of mTBIs, the cellular mechanisms that facilitate or prevent recovery following injury remain poorly defined. Here, we extend our previous work on the role of the protein transglutaminase 2 (TG2) in CNS injury and we hypothesize that transcriptional regulation by TG2 restricts metabolic versatility in astrocytes following TBI, thereby impairing neuronal energetic support and worsening pathological outcomes. We utilized an established weight-drop model of repetitive mTBI followed by multi-parametric analysis of TBI pathology in complete TG2 knockout (TG2-/-) and wild type mice. At 28 days post-injury, TG2-/- mice showed marked attenuation of TBI pathology, compared to wild type mice, in vulnerable white matter and default mode network (DMN) regions, as assessed by diffusion magnetic resonance imaging (MRI), resting-state functional MRI, and immunohistochemistry. Integrated epigenomic, proteomic, and metabolomic profiling of cortical astrocytes isolated 28 days after injury revealed a pronounced metabolic restriction in wild type astrocytes which was remarkably attenuated in the TG2-/- mice. This rescue was associated with a de-repression of gene networks involved in glutamate recycling, lipid metabolism, and metabolic homeostasis. Together, these studies provide novel mechanistic insights into the metabolic dysregulation that characterizes persistent TBI pathology, and establish a foundation for evaluating TG2 as a therapeutic target for TBI.

## Introduction

Traumatic brain injury (TBI) is the most prevalent central nervous system (CNS) injury around the world, with the majority of TBIs classified as mild [1–3]. The primary injury, along with the resulting pathological and symptomatic courses following TBI, is highly heterogeneous, making prognosis difficult to stratify using standard symptom-based assessments and clinical imaging [4, 5]. Even mild TBIs (mTBIs) can result in subsequent reoccurring headaches, disrupted sleep patterns, and cognitive and behavioral impairment, and a large fraction of mTBI patients experience lingering symptoms extending beyond 3 months after injury [6–8]. These injuries also increase the risk of developing neurodegenerative conditions, which is further exacerbated by repetitive injures [9–12]. The long-term pathological and symptomatic sequelae of mTBI are receiving increasing attention, underscoring the need to better understand the evolution of injury processes that underlies chronic pathology.

TBI is multi-factorial disease process that evolves over time and is often characterized by multi-focal axonal injury, neuroinflammation, hemodynamic instability, and metabolic dysregulation [13, 14]. Astrocytes are central to many of these processes, serving as metabolic support hubs and key regulators of CNS injury responses and outcomes [15–18]. Following CNS injury, astrocytes become “reactive”, a term that represents a broad spectrum of morphologic, protein, and metabolic changes that exert both protective and detrimental effects on neurons [19]. In mice, conditional ablation of proliferating reactive astrocytes after a moderate cortical impact TBI results in a dramatic increase in neuronal degeneration and inflammation compared to non-ablated controls, highlighting their protective role [20]. On the other hand, reactive astrocytes can also contribute to injury progression. Subsequent to TBI, astrocytes can exacerbate neuroinflammation [21, 22] and secrete factors that inhibit axonal regeneration and promote synaptic dysfunction [23]. Moreover, metabolic adaptations in astrocytes after a mTBI directly influence their reactive phenotypic state, shifting them toward either neuro-supportive or neurotoxic functions [24]. Understanding the mechanisms that govern these TBI-induced metabolic and physiological changes in astrocytes is critical for developing interventions to mitigate long-term adverse outcomes. One key factor influencing astrocyte responses to CNS injury is Transglutaminase 2 (TG2).

TG2 is a functionally versatile protein expressed across all brain cell types with well-established roles in regulating cell survival and injury responses [25–28]. Its well-characterized activities include calcium–dependent transamidation and protein scaffolding, with the former allowing it to post–translationally modify proteins [29–32]. TG2 expression is induced by injury signaling, and althoughTG2 is predominantly cytosolic, it can shuttle in and out of the nucleus, where it is specifically enriched in euchromatin and it can interact with a wide range of nuclear proteins (e.g. transcription factors, chromatin regulatory complexes) to influence gene expression and cell survival in injury contexts [25, 33–36]. Through a combination of its catalytic and potential scaffolding activity in the nucleus, it can both facilitate and repress gene expression depending on signaling contexts [32, 37–43]. Our previous studies show that TG2 plays a significant role in shaping CNS injury outcomes. Complete TG2 knockout mice exhibited an approximately 36% reduction of infarct volumes compared to wild type mice 24 hours after middle cerebral artery ligation [26]. Similarly, following bilateral contusion spinal cord injury, mice with astrocyte-selective deletion of TG2 showed a significantly faster and greater recovery of motor function compared to controls [44]. *In vitro*, TG2 depletion in primary astrocytes enhances their ability to protect co-cultured neurons in hypoxic conditions and significantly increases their ability to promote neurite outgrowth on an inhibitory matrix [28, 37, 42]. Our previous findings indicate that astrocytic TG2 acts predominantly as a transcriptional repressor under injury conditions, and that the neurosupportive effects of TG2 deletion are likely due in part to altered regulation of astrocytic metabolic pathways, including upregulations in fatty acid transport, uptake, and utilization pathways [42, 44].

We hypothesize that TG2-dependent regulation of astrocytic gene expression contributes to worsened outcomes following CNS injury by promoting a metabolically restrictive phenotype in astrocytes. In this study, we employed a repetitive, mTBI model for its established association with long-term metabolic dysregulation [13, 14, 45, 46] and its compatibility with multi-parametric MRI analyses paired with ex vivo astrocyte profiling. This approach enabled us to assess how TG2 deletion influences long-term astrocyte genetic regulation, function, and pathology in mice following TBI. One month after injury, we identified changes in spontaneous low-frequency oscillations in the blood oxygen level dependent (BOLD) signal, functional connectivity, white matter integrity, and GFAP-dependent astrocyte reactivity largely overlapping with default mode network (DMN) regions. Focused analysis of isolated cortical astrocytes from TBI brains revealed prominent epigenetic inhibition, protein downregulation across metabolic pathways, and metabolomic changes consistent with a metabolic restriction.

## Results

### 1. rmTBI induction using a controlled closed–skull weight drop model

To model repetitive mTBI (rmTBI), we used a weight drop apparatus that was custom-modified to deliver highly reproducible mild closed-skull impacts with linear and angular acceleration (Extended Data Fig. 1a-d), as previously described [47]. The established conditions of this design (e.g. weight, height of sled, impactor tip, and spring) produce consistent, diffuse impact forces, and do not cause skull fractures or hemorrhage. Following our experimental timeline (Extended Data Fig. 1e), cohorts of wild type (WT) and TG2-/- male C57/Bl6N mice, at 10-12 weeks of age, were separated into TBI or sham groups. For MRI experiments, these groups received baseline MRIs, consisting of T2-structural imaging, resting state functional MRI (rs-fMRI) and diffusion MRI (dMRI). Following baseline imaging, mice were given mTBIs (or sham conditions) for 5 consecutive days. We measured the recovery of righting reflex after TBI, a common measure for return of consciousness that varies according to TBI severity and level of anesthesia [48], which was 25.26 ± 0.78 seconds for sham groups (n=14 per genotype) and 35.78 ± 2.86 seconds for TBI groups (n=14 per genotype). No significant difference was found between genotypes. We also used high speed videos to measure head angular accelerations after impacts using kinematic software [49], as described in [47], to track angular displacement of the mouse head over time. We observed a mean maximum downward (clockwise with a rightward facing head) angular acceleration of 4,149 rads/sec, which is consistent with a mild TBI (Extended Data Fig. 1d). These data did not show a significant whiplash effect in these cohorts. Twenty-eight days after the last impact, the mice received post-TBI MRIs, and their brains were harvested for cortical astrocyte isolation (Extended Data Fig. 1e). Parallel groups were also harvested at 28 days post-TBI for whole brain fixation for immunohistochemistry.

### 2. Effects of TG2 and rmTBI on brain amplitude of low frequency fluctuations (ALFFs) in rs-fMRI

ALFF quantifies the fluctuations of BOLD signal in the 0.01–0.1 Hz range, reflecting the spontaneous neural activity in the brain during rest [50]. We generated group-specific ALFF maps to observe changes in the ALFF patterns, indicative of spontaneous neural activity in local brain areas at rest, in each genotype (WT and TG2-/-) in different conditions (baseline, sham, and post-TBI) (Fig. 1). At whole brain scale, baseline and sham TG2-/- mice showed more suppressed ALFF (mean ± SD, baseline: 1.05 ± 0.17 and sham: 1.12 ± 0.08) than WT counterparts (baseline: 1.35 ± 0.11 and sham: 1.37 ± 0.14) (Fig. 1a, b). However, following rmTBI, ALFF in the TG2-/- group indicated greater spontaneous activity (mean ± SD, 1.37 ± 0.14) than WT (mean ± SD, 1.15 ± 0.11), highlighting the role of TG2 in regulating neural activity in response to rmTBI (Fig. 1c). As the rs-fMRI data were co-registered into the mouse brain atlas [51], we next computed mean ALFF from 70 brain regions from each subject. A 2 × 2 ANOVA revealed significant effects of genotype and rmTBI as well as interaction between them on ALFF in 3 brain regions (prelimbic cortex – PL, infralimbic cortex – ILA, and thalamus – TH) and significant interactions between genotype and rmTBI, without individual variable main effects, were identified in 4 regions (motor cortex – MO, somatosensory cortex – SS, retrosplenial cortex – RSP, and hippocampus – HIP) as described below (Fig. 2).

**Figure 1.**
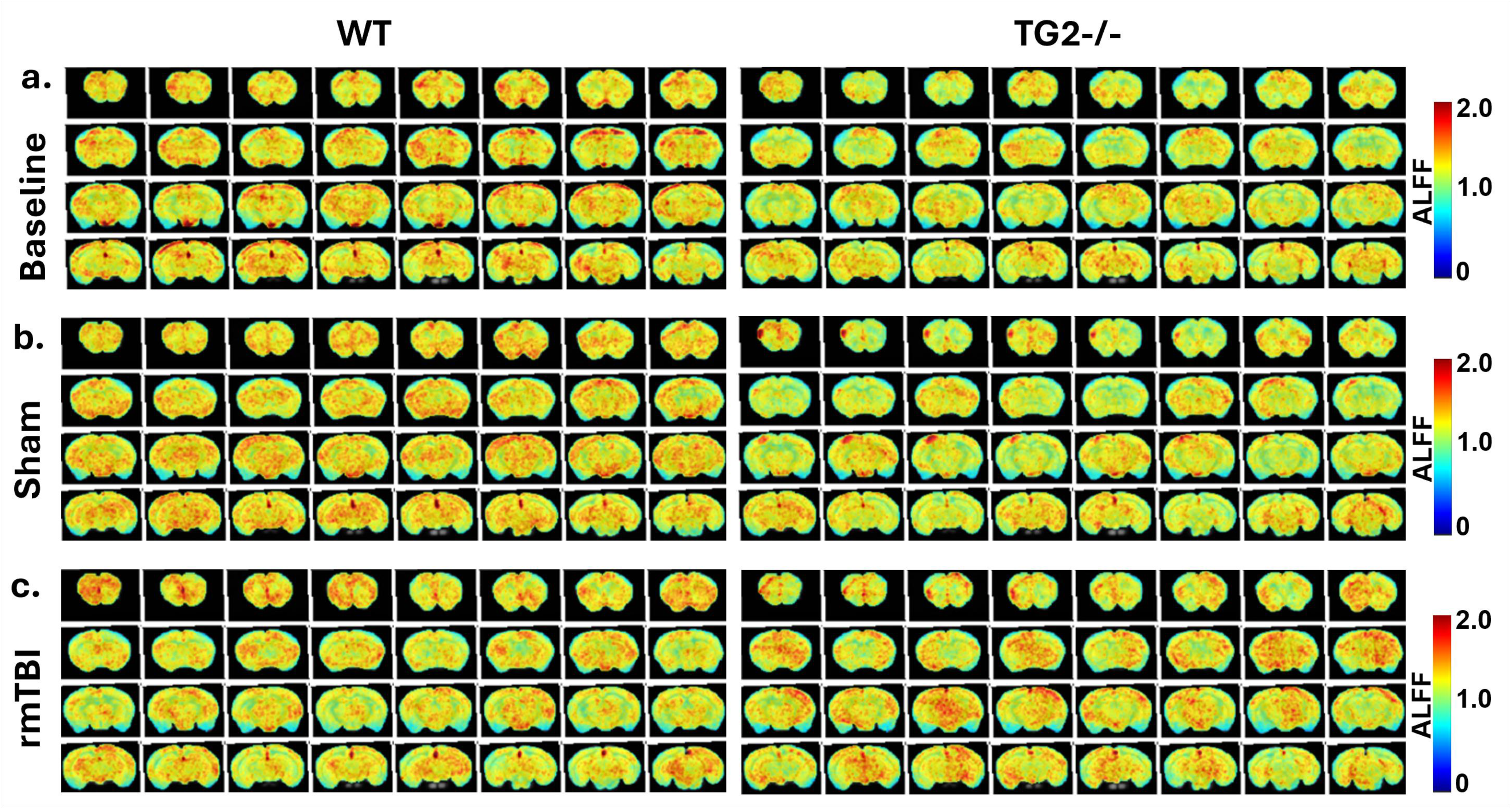
Effects of TG2 and rmTBI on brain amplitude of low frequency fluctuations (ALFFs). ALFF maps of the WT (left) and TG2-/- (right) groups at different conditions: **a)** Baseline, **b)** post-sham, and **c)** post-TBI. Hot and cold colors represent enhanced or suppressed ALFF, respectively. Anatomical regions were identified by mapping onto an in-house mouse brain atlas [51]. (n= 6 mice per group).

**Figure 2.**
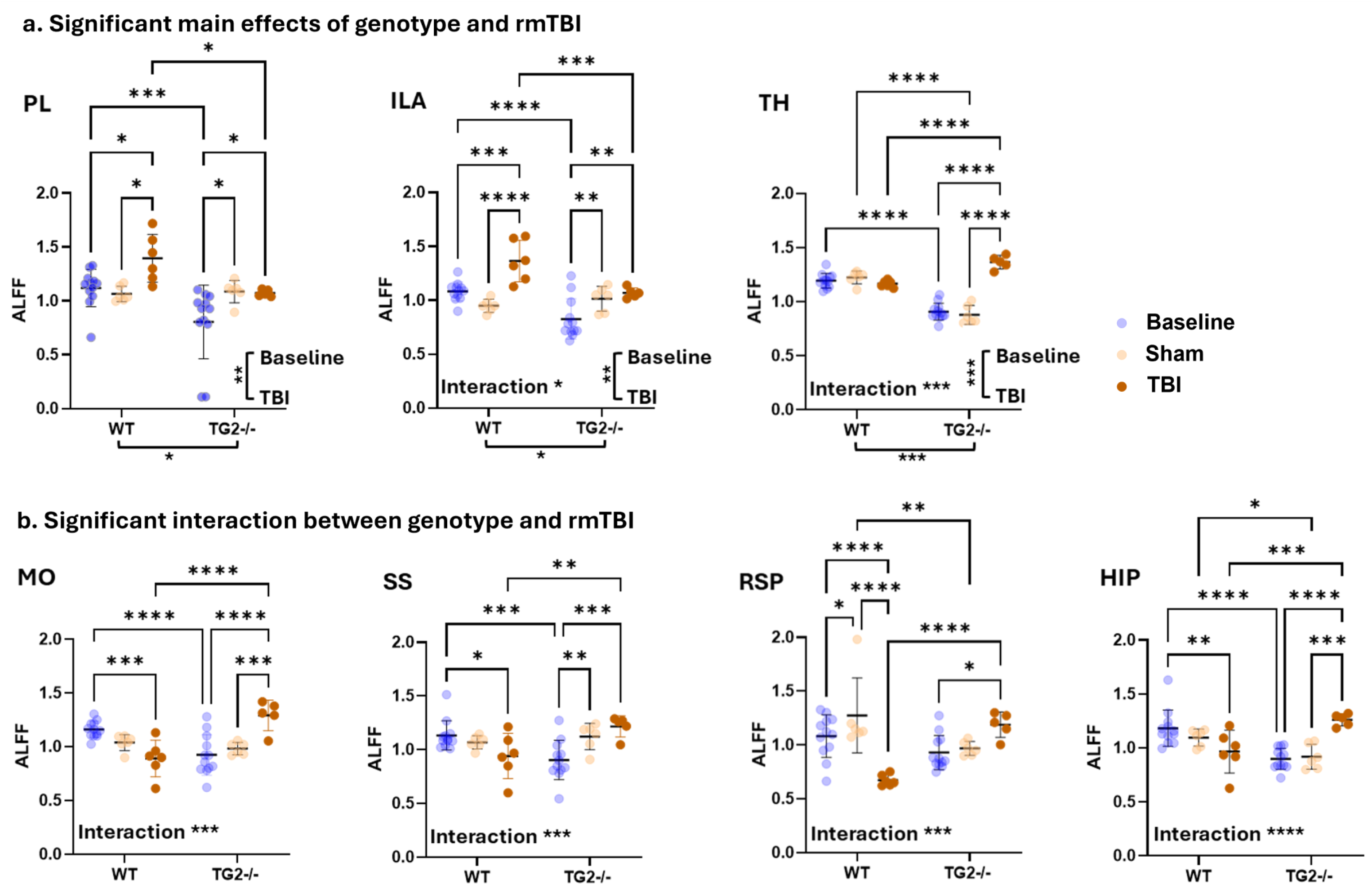
WT and TG2-/- mice show divergent ALFF responses following rmTBI. **a)** Significant main effects of genotype and mTBI on ALFF: prelimbic cortex (PL), infralimbic cortex (IL), and thalamus (TH). **b)** Significant interaction between genotype and mTBI variables: motor cortex (MO), somatosensory cortex (SS), retrosplenial cortex (RSP), and hippocampus (HIP). (n= 6 mice per group, two-way ANOVA with Tukey’s HSD multiple comparison tests, α ≤ 0.05; *P <0.05, **P <0.01, ***P <0.001, and ****P <0.0001.

In the PL (main effects of genotype: F1,41 = 6.34, p = 0.04; rmTBI: F2,41 = 6.0, p = 0.005, and interaction: F2,41 = 2.73, p = 0.05) and ILA (main effects of genotype: F1,41 = 7.1, p = 0.03; rmTBI: F2,41 = 10.86, p = 0.005, and interaction: F2,41 = 2.58, p = 0.05), TBI WT mice had greater ALFF relative to the TBI TG2-/- group (Fig. 2a). In contrast, the TBI TG2-/- group exhibited significant enhancement in thalamic ALFF compared to WT counterparts (main effects of genotype: F1,41 = 36.47, p < 0.001; rmTBI: F2,41 = 31.28, p < 0.001, and interaction: F2,41 = 12.12, p < 0.001) (Fig. 2a).

Significant interactions between genotype and rmTBI, without significant main effects in either variable, were identified in the MO (genotype: F1,41 = 0.77, p = 0.38; rmTBI: F2,41 = 1.06, p = 0.35, and interaction: F2,41 = 12.30, p < 0.001), SS (genotype: F1,41 = 0.52, p = .48; rmTBI: F2,41 = 1.2, p = 0.30, and interaction: F2,41 = 11.26, p < 0.001), RSP (genotype: F1,41 = .10, p = 0.74; rmTBI: F2,41 = 3.1, p = 0.06, and interaction: F2,41 = 16.41, p < 0.0001), and HIP (genotype: F1,41 = 1.80, p = 0.18; rmTBI: F2,41 = 1.9, p = 0.15, and interaction: F2,41 = 18.20, p < 0.0001) (Fig. 2b).

### 3. Effects of genotype and rmTBI on resting-state functional connectivity (rsFC)

A representative image of the 21 brain regions used for connectivity analysis is shown in Figure 3a (black circles), and a representative connectivity network where nodes are interconnected via edges (black lines) is shown in Figure 3b. We observed a significant genotype effect (2 × 2 ANOVA) on PL-CA1, PL-CA3, and ILA-CA2 rsFC of the cortico-hippocampal network (Fig. 3c and Extended Data Fig. 2). Tukey’s post-hoc analysis specifically showed genotype rsFC differences after rmTBI (PL-CA1: p < 0.001, Cohen’s d = −0.42; PL-CA3: p = 0.0026, Cohen’s d = −0.31; ILA – CA2: p = 0.002, Cohen’s d = −0.32), but not at baseline (PL-CA1: p = 0.69, Cohen’s d = −0.02; PL-CA3: p = 0.53, Cohen’s d = 0.03; ILA-CA2: p = 0.07, Cohen’s d = −0.1) or in sham conditions (PL-CA1: p = 0.95, Cohen’s d = −0.003; PL-CA3: p = 0.28, Cohen’s d = −0.07; ILA-CA2: p = 0.07, Cohen’s d = −0.08) (Fig. 3c).

**Figure 3.**
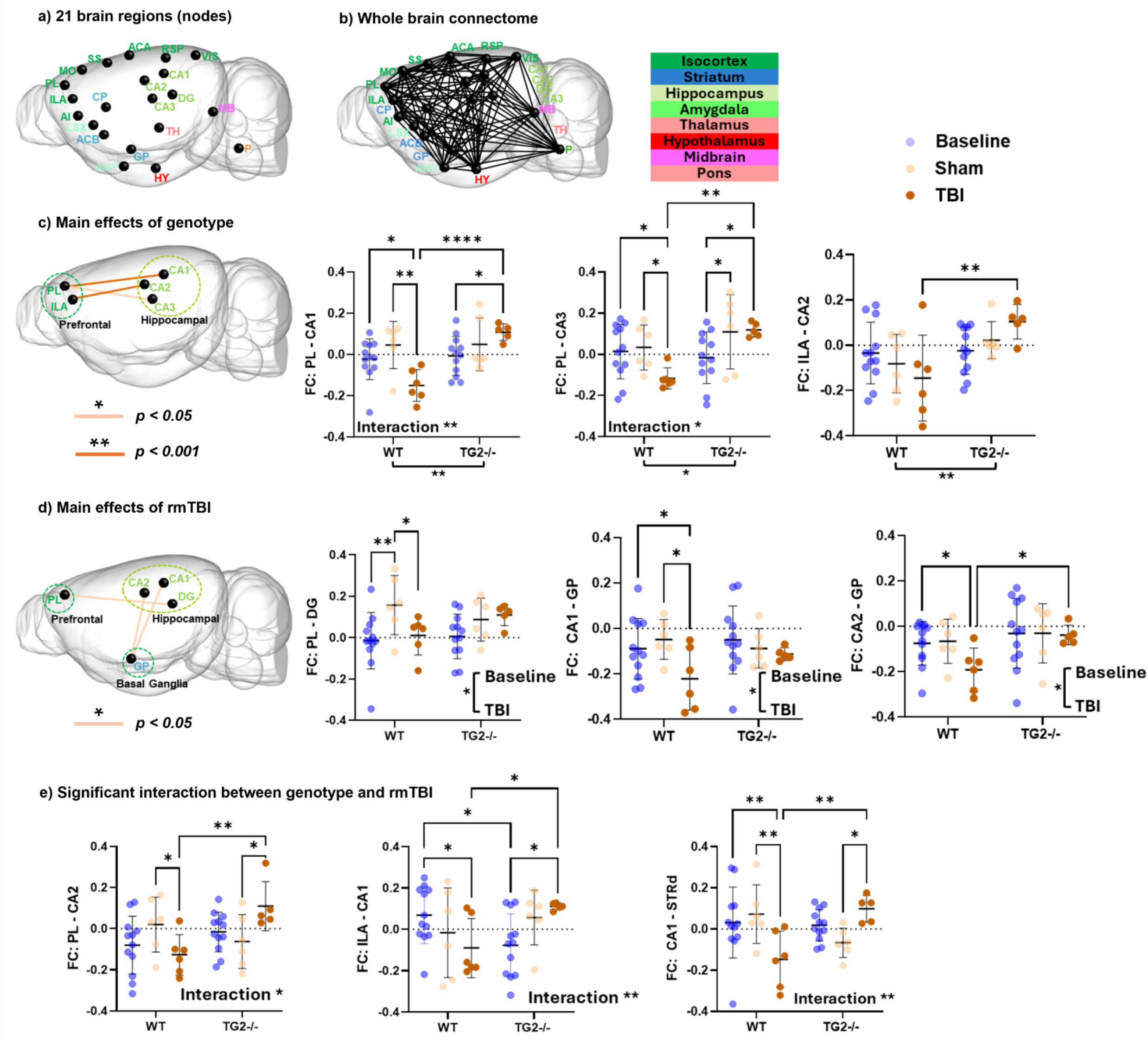
Functional connectivity (FC) changes across WT brain regions after rmTBI are largely reversed in TG2-/- mice. **a)** 21 brain regions (black nodes) included in the node-to-node connectivity analysis, **b)** whole brain connectivity network among the 21 nodes. Black lines represent the connection between the nodes, **c)** FC between brain regions where a significant main effect of genotype was observed, **d)** FC that was significantly altered by rmTBI, and **e)** significant genotype × rmTBI interaction. Color of the line represents the level of significance as denoted in the figure. (n= 6 mice per group, two-way ANOVA with Tukey’s HSD multiple comparison tests, α ≤0.05. *P <0.05, **P <0.01, ***P <0.001).

We observed a significant effect of rmTBI (2 × 2 ANOVA) on PL-DG rsFC of the cortico-hippocampal network, and CA1-GP and CA2-GP of the hippocampal-basal ganglia network (Fig. 3d and Extended Data Fig. 2). Tukey’s post-hoc analysis showed rmTBI rsFC differences in the WT group (PL-DG: p = 0.02, Cohen’s d = 0.21; CA1-GP: p = 0.02, Cohen’s d = 0.22; CA2 – GP: p = 0.05, Cohen’s d = 0.19), but not in the TG2-/-group (PL-DG: p = 0.75, Cohen’s d = −0.02; CA1-GP: p = 0.73, Cohen’s d = 0.02; CA2-GP: p = 0.91, Cohen’s d = 0.008) (Fig. 3d). In addition, significant interaction between genotype and rmTBI (2 × 2 ANOVA) was observed in the following pathways: PL-CA2 (F2,41 = 5.0, p = 0.01), PL-CA2 (F2,41 = 5.0, p = 0.01), ILA-CA1 (F2,41 = 5.85, p = 0.005), and CA1-STRd (F2,41 = 7.22, p = 0.002) (Fig. 3e). In all identified pathways (Fig. 3c-e), functional connection strength was significantly reduced or trended towards a decrease in TBI WT mice relative to WT sham, whereas TG2–/– mice showed no significant differences or trended towards an increase in connection strength in TBI groups relative to sham.

### 4. Effects of genotype and rmTBI on brain Fractional Anisotropy (FA)

We examined the effects of genotype and rmTBI on brain white matter integrity by measuring the overall degree of water diffusivity, i.e. FA, along the major white matter bundles, like corpus callosum (cc), corticospinal tract (cst), and stria terminalis (st). We observed that WT mice exhibit decreased FA (Fig. 4a) whereas TG2-/- mice tend to preserve the white matter integrity (Fig. 4b) following rmTBI. A 2 × 2 ANOVA revealed significant effects of genotype, rmTBI, and interaction between these variables in three major white matter bundles as described below:

**Figure 4.**
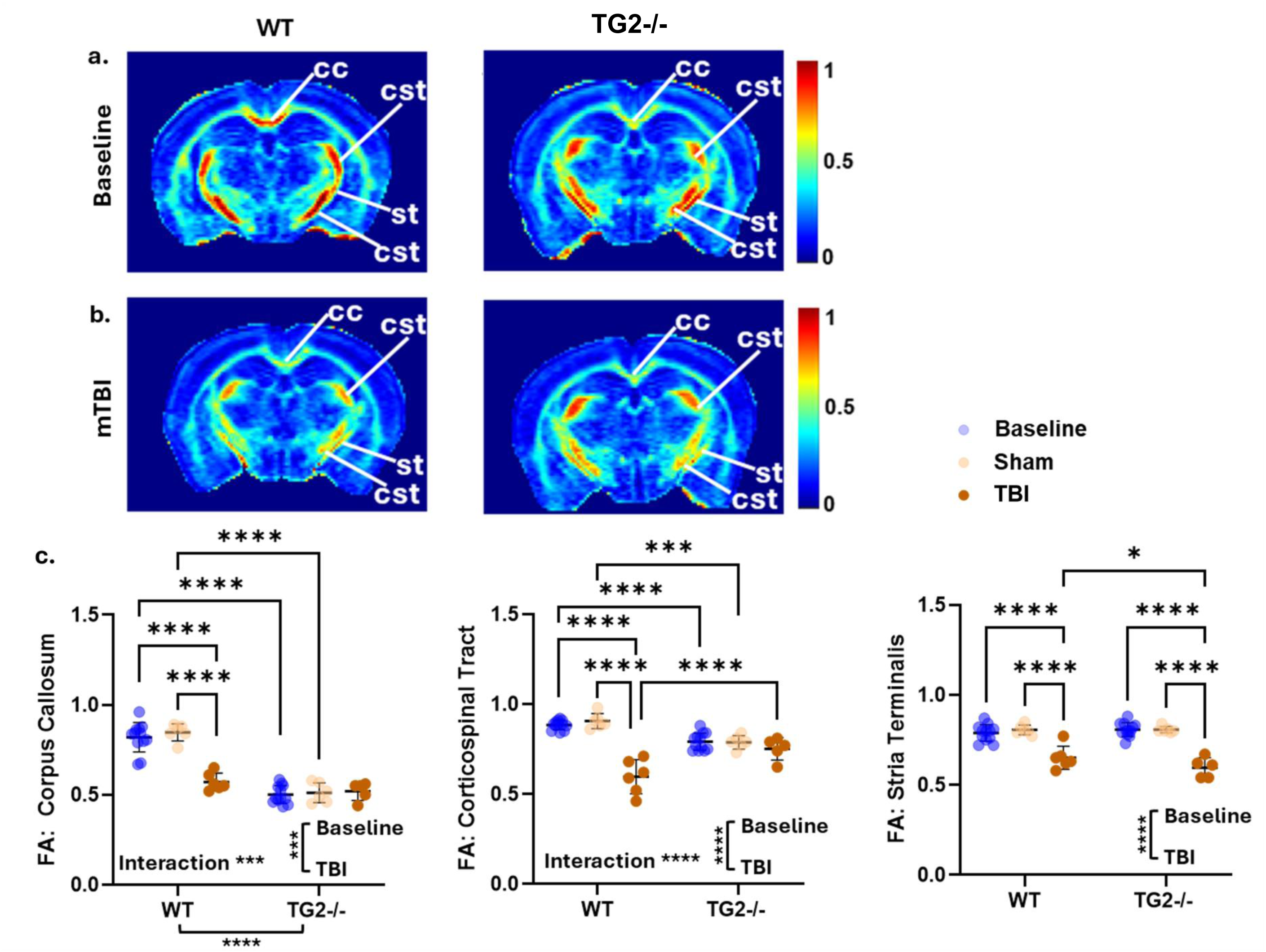
TG2-/- mice show greater preservation of white matter tract integrity after rmTBI compared to WTs. Fractional anisotropy (FA) maps represent white matter microstructural integrity in the WT and TG2-/-groups at **a)** baseline and **b)** following rmTBI, and **c)** quantification of the mean FA showing significant effects of rmTBI in three major WM bundles: corpus callosum (cc), corticospinal tract (cst), and stria terminalis (st). Higher FA indicates greater microstructural integrity and vice-versa. (n= 6 mice per group, two-way ANOVA with Holm-Šidák multiple comparison tests, α ≤0.05; *P < 0.05, ****P <0.0001).

In CC, main effects of genotype (F1,41 = 156.7, p < 0.0001), rmTBI (F2,41 = 17.18, p < 0.0001), and interaction (F2,41 = 21.24, p < 0.0001) were observed. Tukey’s post-hoc analysis showed a significant decrease in FA in the WT group after rmTBI (p < 0.0001, Cohen’s d = 0.66), but not in the TG2-/- group (p = 0.58, Cohen’s d = −0.04) (Fig. 4c).

In the cst, we did not observe any effects of genotype (F1,41 = 1.34, p = 0.25), but we found effects of rmTBI (F2,41 = 44.06, p < 0.0001) and the interaction (F2,41 = 26.21, p < 0.0001) between them. Like CC, FA in the cst was significantly decreased in the WT group after rmTBI (p < 0.0001, Cohen’s d = 0.98), but not in the TG2-/- group (p = 0.25, Cohen’s d = 0.09) (Fig. 4c).

In st, we observed a significant main effect of rmTBI on FA (F2,41 = 68.37, p < 0.0001), but not of genotype (F1,41 = 0.77, p = 0.39), or of the interaction between them (F2,41 = 2.70, p = 0.07). Unlike CC and cst, post-hoc analysis revealed significant reduction of FA in both WT (p < 0.0001, Cohen’s d = 0.57) and TG2-/- (p < 0.0001, Cohen’s d = 0.80) groups after rmTBI (Fig. 4c).

### 5. TG2 deletion attenuates GFAP-dependent reactive gliosis following rmTBI

Reactive astrogliosis with GFAP upregulation is commonly observed across TBI models and the GFAP blood biomarker is one of the most reliable clinical indicators of TBI severity [5, 14, 52–54]. Therefore, we assessed GFAP expression across multiple brain regions one month after rmTBI using immunofluorescence. In low magnification surveys across brain regions, the TBI WT group showed prominent increases in GFAP expression in white matter tracts, specifically the CC and cingulate white matter bundle (CWMB), as well as increased levels in HIP and the RSP, a hub of the DMN and a region underlying the site of impact (Fig. 5a). We selected the CC, CWMB, RSP, and ventral RSP (vRSP) for quantification of GFAP levels at higher magnification (Fig. 5b). Compared to sham, the TBI WT group showed significant increases in the area fraction of GFAP expression in the CWMB, RSP, and vRSP, with an upward trend in CC. Remarkably, the TBI TG2-/- group showed no significant increases in GFAP levels compared to sham across the regions analyzed, however there was an upward trend in the RSP (Fig. 5c-g). Sham WT and TG2-/- groups showed similar levels of GFAP across brain regions (Fig. 5a, d-g). We also analyzed NeuN immunofluorescence as an indicator for neuronal viability within our groups. NeuN is a well-known neuronal nuclear marker and RNA-splicing regulator that shows decreased or mislocalized expression in injury states in association with neuronal stress [55]. TBI WT mice showed a consistent significant decrease in NeuN immunoreactivity compared to sham WT across all regions analyzed, but the TBI TG2-/- showed an attenuated decrease that was not significantly different from sham levels (Extended Data Fig. 3).

**Figure 5.**
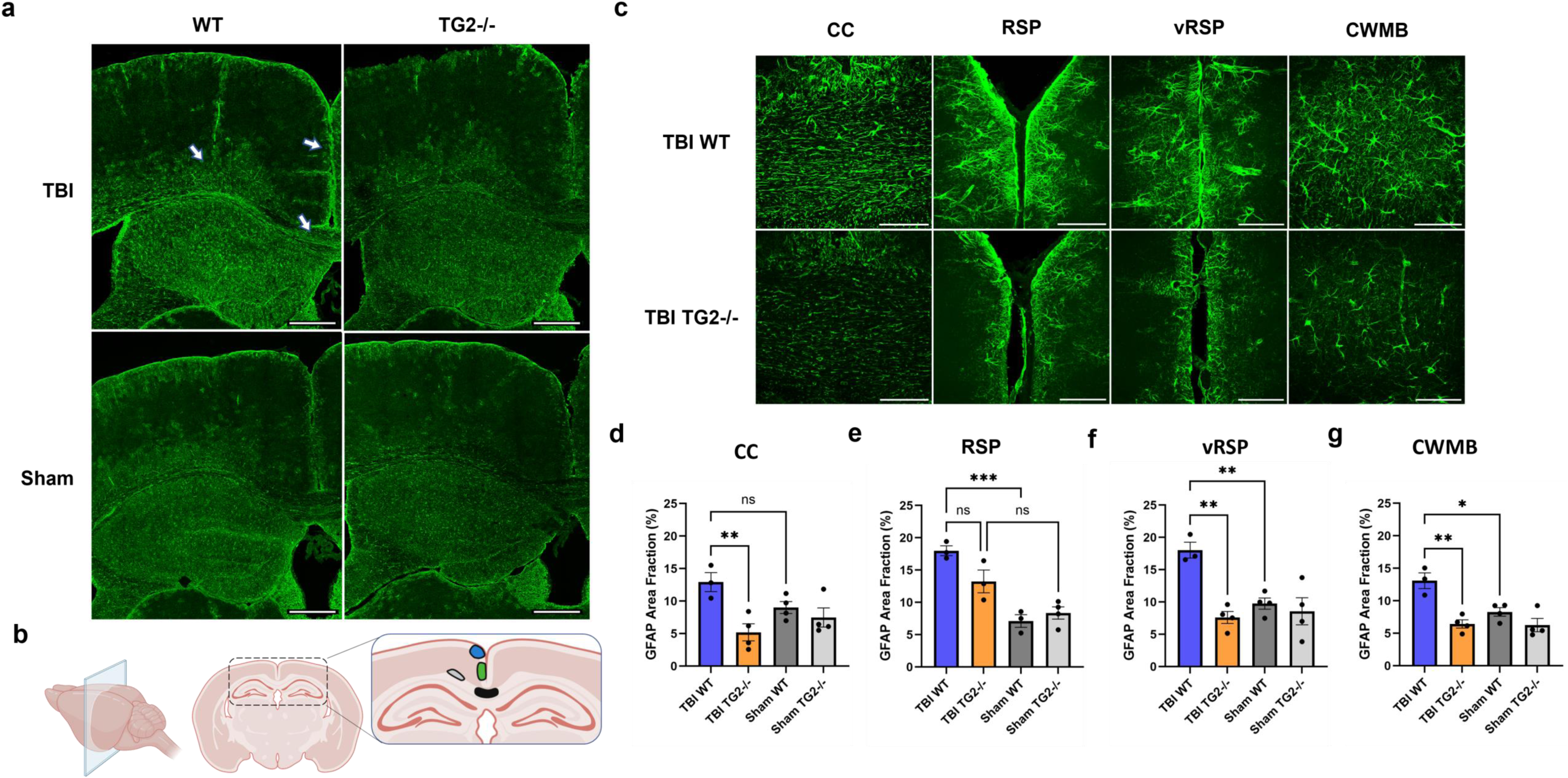
**TG2-/- mice show attenuated reactive gliosis following rmTBI**. **a)** Representative 4x images of GFAP immunoreactivity in each group. White arrows in TBI WT group show increased GFAP labeling in vRSP (ventral retrosplenial cortex), CWMB (cingulate white matter bundle), and CC (corpus callosum) scale bar = 400 μm. **b)** Diagram of regions within coronal sections selected for GFAP quantification (colored areas); blue– RSP (retrosplenial cortex), green– vRSP, black– CC, gray– CWMB. **c)** Representative 40x images (scale bar = 70 μm) and **(d-g)** quantifications of GFAP immunoreactivity in CC, RSP, vRSP, and CWMB across groups. Data in bar graphs represent mean ± s.e.m. (n= 3-4 mice per group, two-way ANOVA with Šidák’s multiple comparison test, α ≤ 0.05. *P < 0.05, **P < 0.01, ***P < 0.001, ns, not significant). Diagram created in BioRender.com.

### 6. TG2 deletion reverses injury–induced downregulation of metabolic gene programs in astrocytes

#### TBI-induced changes in astrocyte histone acetylation

We hypothesize that TG2 regulation of gene expression through its nuclear interactions plays an important role in regulating astrocyte reactive phenotype and metabolism following injury. Therefore, we harvested and dissected cortices from the brains of the MRI cohorts and used MACS with anti-ACSA-2 microbeads to isolate cortical astrocytes for epigenetic, proteomic, and metabolomic profiling (Fig. 6a). We previously found that TG2 interacts with Sin3a, a transcriptional corepressor complex that facilitates histone deacetylation by HDAC1/2 and an important regulator of injury response [37, 42, 56, 57]. Therefore, we measured genome wide locations and levels of H3K9 acetylation (H3K9ac), a histone marker of transcriptionally active promoters [58], in cortical astrocytes from sham and TBI brains using CUT&RUN-seq. Following CUT&RUN, we performed paired end sequencing and used MACS2 for peak calling of aligned reads.

**Figure 6.**
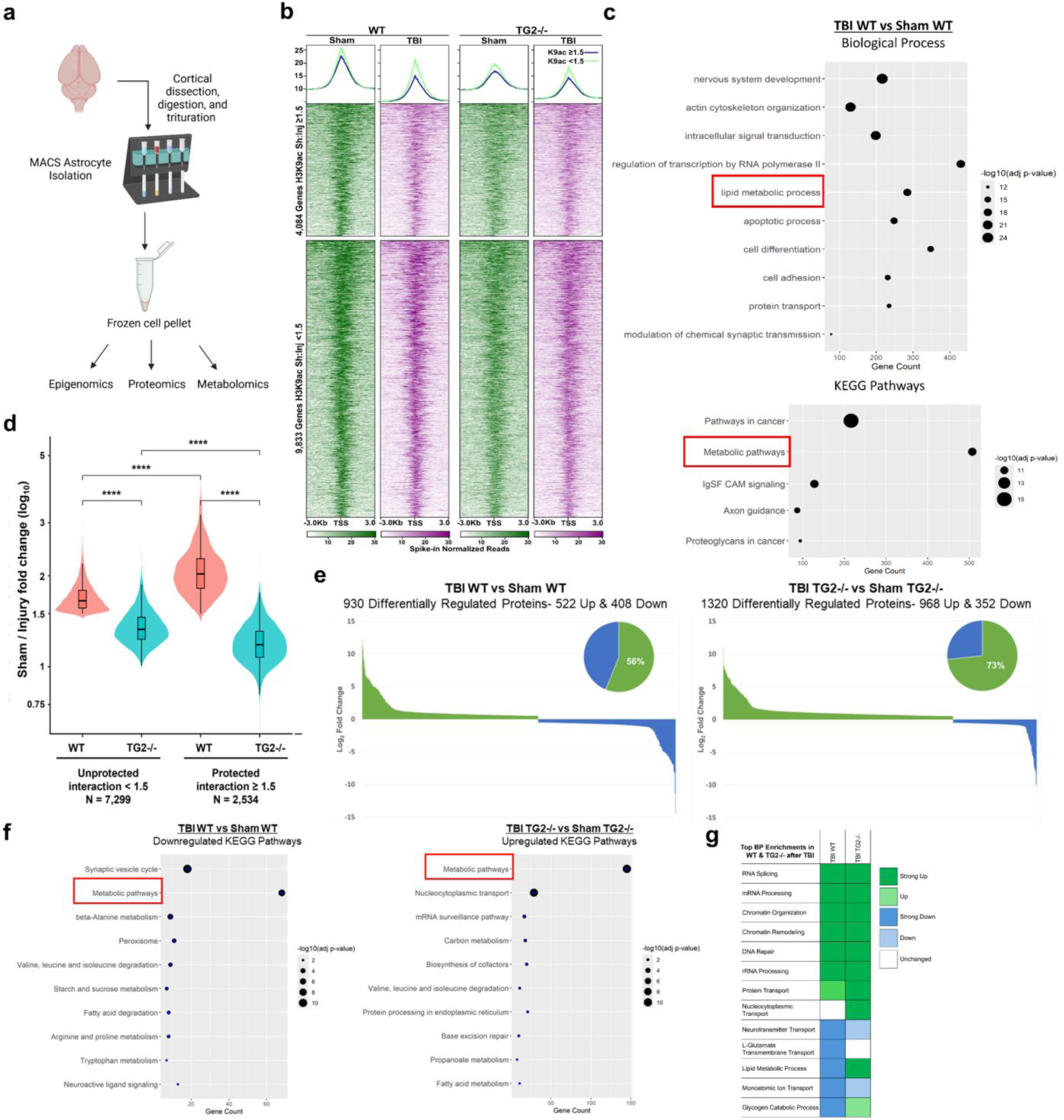
Epigenomic and proteomic profiling of *ex-vivo* astrocytes. **a)** Diagram of cortical astrocyte isolation from TBI and sham brains using MACS with anti-ASCA2 micro magnetic beads. **b)** CUT&RUN-seq enrichment for H3K9ac from cortical astrocyte nuclei from each genotype in TBI and sham groups. Heatmaps show TSS-centered H3K9ac signal sorted by genes with ≥1.5-fold reduction (top) and <1.5-fold reduction (bottom) in H3K9ac comparing Sham WT to TBI WT. **c)** DAVID GO Biological Process and KEGG enrichments for total genes with ≥1.5-fold reduction in WTs following rmTBI. **d)** Violin plots showing Sham vs TBI log_2_ fold change in H3K9ac signal for genes with ≥1.5-fold reduction in H3K9ac signal in both WT and TG2-/- groups after rmTBI (“unprotected”) and for genes with <1.5-fold change only in the TG2-/- group (“protected”). **e)** profiles of up- and down-regulated proteins after rmTBI in each genotype. **f)** KEGG enrichment of <u>downregulated</u> proteins in WTs after rmTBI (left), and <u>upregulated</u> proteins in TG2-/- after rmTBI (right). **g)** Heat map of up- and downregulated pathways in the top GO Biological Process enrichments in WT and TG2-/- astrocytes after TBI. Diagram in **(a)** created in BioRender.com. (a-d, n= 2 mice per group, ****P < 0.0001, e- g, n=6 mice per group).

Almost all H3K9ac peaks (about 30k peaks genome wide) were reduced in WT astrocytes at 28 days post rmTBI, with 9,515 peaks showing a ≥1.5-fold reduction (not shown). Plotting H3K9ac signal at H3K9ac+ transcription start sites (TSS) illustrates this finding, with 4,084 genes showing ≥1.5-fold loss (Fig. 6b). TG2-/- astrocytes showed a similar trend after injury, however the fold-reduction was notably attenuated. Interestingly, the TG2-/- sham profile suggests that there are also baseline differences in H3K9ac distribution compared to WT sham, with the TG2-/- group showing less concentrated TSS signal. The TBI TG2-/- group also retains higher H3K9ac signal in TSS flanking regions compared to the TBI WT group, which may be associated with enhancer sites.

Gene enrichment analysis was performed on all known overlapping genes with ≥1.5-fold loss of H3K9ac (within TSS and gene bodies) in TBI WT compared to sham WT groups. GO Biological Process enrichment showed top enrichment categories which included nervous system development, actin cytoskeleton and regulation of RNA polymerase II. Notably, lipid metabolic process was among these top categories, and KEGG enrichment analysis revealed metabolic pathways as a top category showing the greatest number of enriched genes (Fig. 6c).

Further analysis revealed that among H3K9ac peaks with ≥1.5-fold signal loss in WT after injury (n = 9,833), 2,534 (25.8%) showed a substantially attenuated decrease in TG2−/− astrocytes (interaction ratio ≥ 1.5; median Sham/Injury fold change 1.18 in KO vs 2.03 in WT), indicating protection from injury-induced H3K9ac loss (Fig. 6d). Of the associated genes protected from H3K9ac loss, ∼1/3rd of them are located at TSS and 2/3rd within gene bodies and may be enhancers (not shown). Biological Process enrichment of these protected genes showed top enrichments in positive regulation of RNA polymerase II and nervous system development, similar to enrichments discussed above, with a significant but relatively less prominent metabolic signature focused on lipid metabolism (not shown). Additionally, cell type enrichment analysis of all genes with ≥1.5- fold loss in H3K9ac signal showed a strong astrocyte enrichment, which demonstrates the specificity of our astrocyte isolation (Extended Data Fig. 4).

#### TBI-induced changes in astrocyte proteome

We next analyzed proteomic changes across our astrocyte groups. WT astrocytes showed similar amounts of significantly up- and down-regulated proteins after TBI, while TG2-/- astrocytes showed a predominant upregulation of proteins (Fig. 6e), a profile consistent with TG2 depletion in our previous RNA-seq and proteomic data [28, 42, 44]. Kegg enrichment analysis of differentially regulated proteins showed a prominent downregulation of metabolic pathway proteins in the TBI WT group, including downregulated fatty acid metabolism and branched-chain amino acid breakdown. Strikingly, these categories showed opposite changes in the TBI TG2-/- group, being significantly enriched among upregulated proteins (Fig. 6f). Biological Process enrichment of up- and down-regulated proteins across TBI groups revealed that both WTs and TG2-/- astrocytes exhibit many similar proteomic changes following injury (e.g. RNA splicing, chromatin organization), however the TG2-/- group showed unique regulation of glutamate transport, lipid metabolism, nucleocytoplasmic transport, and glycogen metabolism (Fig. 6g).

We further assessed how H3K9ac gene changes may have contributed to these observed proteomic differences. Biological Process gene enrichment analysis of shared gene IDs between CUT&RUN and proteomic datasets within the TBI WT group showed that proteins involved in prominently downregulated metabolic pathways in Fig. 6g also exhibited ≥1.5-fold H3K9ac reduction at their gene loci, associated with transcriptional repression (Extended Data Fig. 5a). Similar analysis using TG2-/- protected H3K9ac genes overlapped with TBI WT protein changes identified glutamate transport genes as a protected group in TG2-/- animals (Extended Data Fig. 5b). While Extended Data Fig. 5a and the proteomic data in Fig. 6g would predict that this TG2-/- de-repressive effect at H3K9ac loci should also extend at least to glycogen metabolic genes, dynamic H3K9ac deposition and removal according to cellular signaling context may ultimately lead to an average de-repression of a wider set of metabolic genes which we may not capture in one timepoint for CUT&RUN but which are more stably expressed in the proteome. These data further suggest that epigenetic mechanisms beyond H3K9ac regulation more substantially contribute to the striking lipid-metabolism protein upregulation in the TBI TG2-/- astrocytes in Fig. 6g.

### 7. TG2 deletion upregulates pathways associated with metabolic versatility following rmTBI

Our proteomic data shows that after rmTBI, WT astrocytes undergo a multi-pathway metabolic restriction (Fig. 7a), consistent with previous studies showing chronic mitochondrial dysfunction and lipid dysregulation after TBI [14, 45, 59]. This effect is reversed in the TBI TG2-/- group, with a large fraction of upregulated metabolic proteins contributing to TCA anaplerosis and mitochondrial function (Fig. 7b). To investigate the impact of these protein expression changes on the metabolic profiles across our astrocyte groups, we performed LC-MS untargeted metabolomic analysis on isolated cortical astrocytes. From a reference library of 116 common metabolites, we could identify 81 metabolites in our samples for relative abundance analysis. Following injury, WT astrocytes showed decreases in ATP and TCA cycle metabolite abundances (Fig. 7c). Notably, this was coincident with a downregulation of proteins involved in glutamate recycling from synapses (e.g. GLAST and EEAT2, or GLT-1) (Fig. 7a). While glutamate taken up by astrocytes can be recycled back to neurons through glutamine, a substantial portion of this glutamate is funneled into mitochondria for oxidation to restore TCA-related intermediates like aspartate. Through this TCA route, glutamate may also contribute to lactate production, potentially via phosphoenolpyruvate carboxykinase and/or malic enzyme [60, 61], which is exported to neurons for energetic support while simultaneously controlling glutamate excitotoxicity. The TBI WT profile indicative of decreased uptake and utilization of synaptic glutamate is therefore consistent with reduced TCA flux and consequent neuronal energetic and excitotoxic dysregulation (Fig. 7c). Among the few upregulated metabolites in the TBI WT group, increased fructose-1,6-bisphosphate and gluconate indicate increased glycolytic flux (Fig. 7c).

**Figure 7.**
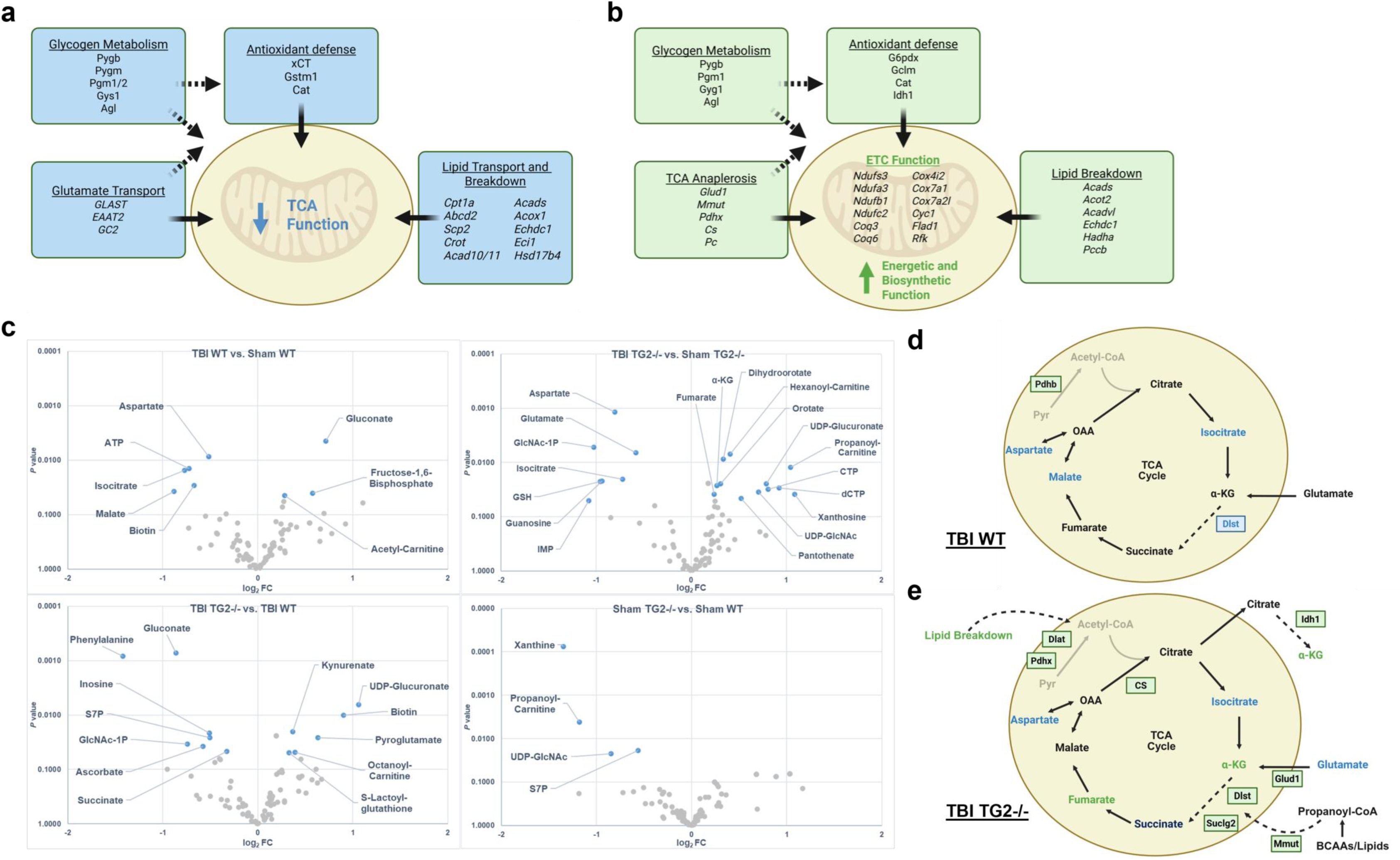
Integrated proteomic and metabolomic profiling of *ex-vivo* astrocytes. Metabolic pathway genes from proteomic KEGG enrichments (Figure 6f) related to TCA and mitochondrial function were summarized for **a)** TBI WT and **b)** TBI TG2-/- groups. **c)** LC-MS metabolomic analysis of isolated cortical astrocytes showing metabolite relative abundance changes for all group comparisons, plotted by p value and log2 fold change. Metabolites that were significantly changed (P < 0.05, pairwise t-tests) are shaded in blue and labeled (n= 8 mice per group). Pathway summaries show differential regulation of TCA-related proteins and metabolites in the **d)** TBI WT and **e)** TBI TG2-/- groups. Diagrams created in BioRender.com.

Contrary to the WTs, TG2-/- astrocytes did not show a complete downregulation of TCA metabolite abundance. While there was a similar decrease in aspartate and isocitrate, they maintained stable levels of malate and increased levels of α-ketoglutarate and fumarate (Fig. 7c). Integrating these findings with our overall proteomic data suggests that these increases are at least partly attributed to an upregulation of anaplerotic pathways after injury (including glutamate, branched-chain amino acid, and lipid metabolism), which does not occur in WTs (Fig. 7d,e). Additionally, although aspartate was decreased in TG2-/- astrocytes after injury, higher abundances of metabolites involved in and derived from pyrimidine synthesis (dihydroorotate, orotate, dCTP, CTP, UDP-GlcNAc) suggest that this group also has a greater utilization of aspartate and likely maintains greater aspartate availability, as it is a necessary precursor in this pathway (Fig. 7c). Similarly, decreased glutamate levels in this group may be a result of greater utilization in TCA cycle (Glud1 upregulation) and glutathione synthesis (Gclm upregulation) (Fig. 7b,e). Overall, TG2 deletion promotes adaptive metabolic gene and protein expression in astrocytes that facilitate neuronal support following injury; these mechanisms are summarized in Fig. 8.

**Figure 8.**
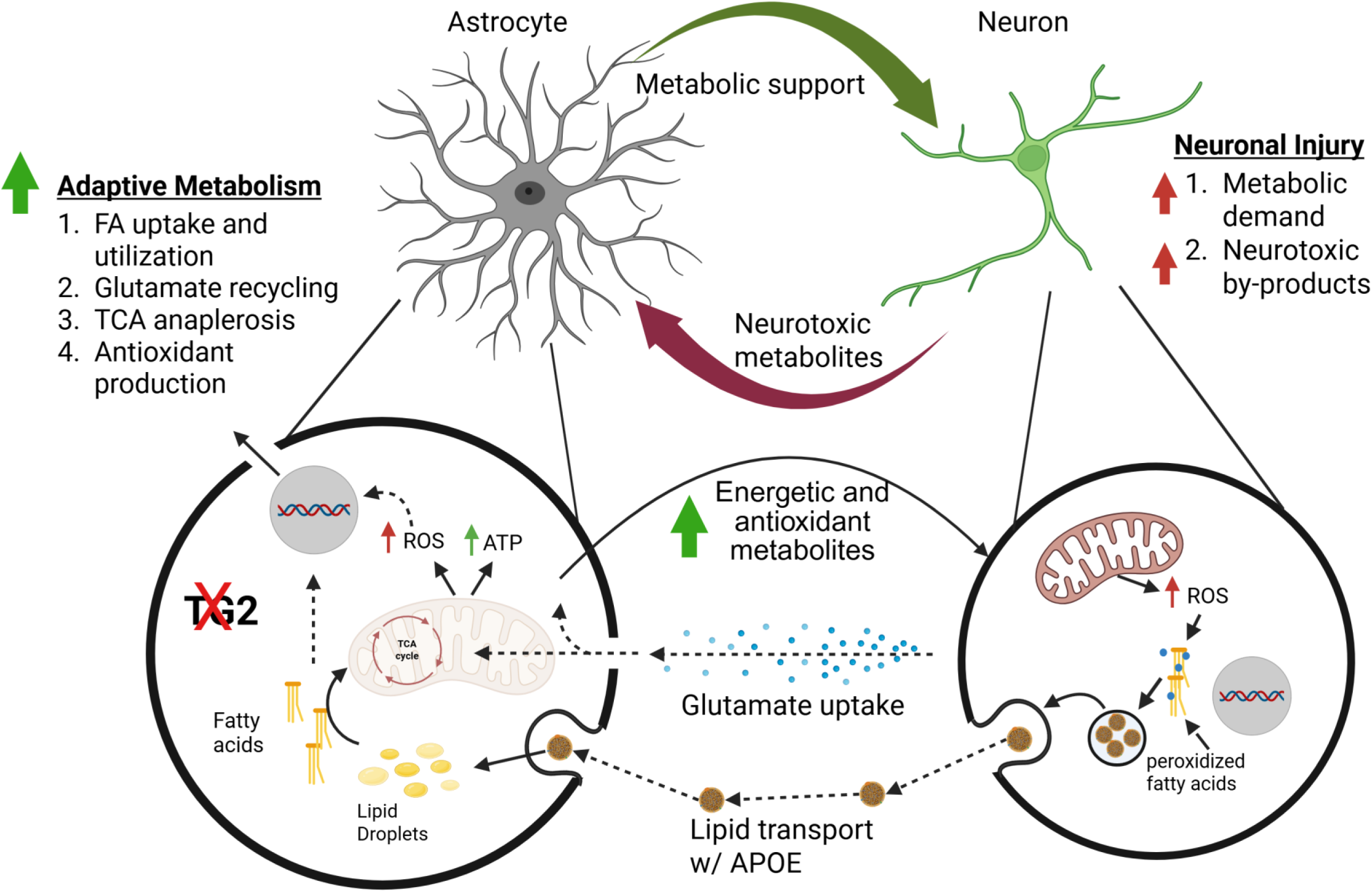
Summary of TG2-regulated astrocytic metabolic adaptations that shape neuronal recovery after rmTBI. TBI and associated axonal injuries disrupt ionic homeostasis, synaptic signaling, and axonal transport, triggering oxidative stress and glutamate-dependent excitotoxicity. These disruptions impose a substantial energetic burden on neurons as they re-establish ion gradients and initiate repair. Although astrocytes are tightly coupled to neuronal energetic needs, subacute rmTBI induces a reactive state in wild-type astrocytes characterized by pronounced epigenetic repression and metabolic restriction. In contrast, TG2-/- astrocytes upregulate pathways involved in lipid transport and utilization, antioxidant production, and TCA cycle anaplerosis, and they maintain stable expression of proteins involved in glutamate transport, overall conferring greater metabolic flexibility to support injured neurons. Created in BioRender.com.

### 8. TG2-/- astrocytes better support neuronal redox balance *in vitro*

Given our findings that TG2-/- astrocytes maintain greater metabolic flexibility in injury conditions, we traced our steps back to determine whether our updated mechanistic model was supported by our previous *in vitro* studies. We have shown in neuron-astrocyte co-cultures that astrocytes with TG2 depletion better support neurite outgrowth on a chondroitin sulfate proteoglycan (CSPG) growth-inhibitory matrix [37, 42]. CSPGs are well known regulators of CNS repair and regeneration after injury, and they have been shown to inhibit autophagic flux in neurons [62, 63]. Therefore, we asked: (1) whether CSPGs induce energetic and oxidative stress in neurons, and if so, (2) whether TG2-/- astrocytes can better protect neurons against these stressors through metabolic support compared to WTs. Indeed, using immunofluorescence to probe for 8-oxo-guanine, the most prevalent marker of DNA oxidation, we found that neurons cultured on CSPGs with poly-D-lysine (PDL) exhibit increased oxidative stress compared to those cultured on PDL alone (Extended Data Fig. 6a,b).

To assess the impact of astrocytic TG2, we co-cultured WT and TG2-/- astrocytes at 3-4 days in vitro (DIV) with neurons grown on either CSPG+PDL or PDL matrices (DIV 9) for 5 days (to DIV 14) using a coverslip-paraffin sandwich design (Extended Data Fig. 6c) [64]. As expected, the addition of either astrocyte group reduced neuronal oxidative stress on both growth matrices (Extended Data Fig. 6b,e). In PDL groups, compared to WTs, TG2-/- astrocytes exhibited lower oxidative stress and they better protected neurons against oxidative stress on the PDL matrix (Extended Data Fig. 6d,e). However. in the CSPG condition, there was no difference between neuron pairs, and TG2-/- astrocytes intriguingly exhibited increased oxidative stress (Extended Data Fig. 6f,g). This may be explained by the observation that CSPG neuron cultures paired with TG2-/- astrocytes exhibited increased network density by DIV 14 relative to WT pairs (not shown), consistent with our previous CSPG neurite outgrowth data from 4-day co-culture (neuron DIV 1-DIV 5) [37]; therefore, potential differences in energetic demand due to network complexity may affect neuronal and astrocytic oxidative stress in the CSPG condition.

### 9. TG2-/- astrocytes exhibit increased oxidative stress and mitochondrial fragmentation in association with lipid utilization

Astrocytic TG2 depletion is associated with prominent RNA and protein upregulations in lipid metabolic pathways, and we previously postulated that this contributes to enhanced neuronal metabolic support [42, 44]. To probe for differences in lipid utilization, we assessed oxidative stress and mitochondrial morphology in our astrocyte cultures across different nutrient contexts, with or without palmitate (C16), as both reactive oxygen species generation and mitochondrial fragmentation are indicative of lipid catabolism [65–67]. In high glucose conditions (HG; ∼33 mM), incubation with high concentrations of lipid (50 μM) for 24 hours increased oxidative stress in TG2-/-, but not WT astrocytes, indicating greater lipid catabolism (Extended Data Fig. 7a,b). Mitochondrial morphology analysis under matched conditions showed increased fragmentation in TG2–/– astrocytes irrespective of media condition (Extended Data Fig. 7c,d). Given that mitochondrial fragmentation facilitates CPT1a–dependent fatty–acid import and oxidation [65], this phenotype further indicates elevated lipid utilization in TG2–/– astrocytes.

1. 10. Metabolomics and isotope tracing of neuron-astrocyte co-cultures

We applied a similar co-culture design as above to analyze the metabolome of WT and TG2-/- astrocyte-neuron pairs. WT and TG2-/- astrocytes were incubated with uniformly labeled ^13^C palmitate (U-^13^C C16) for 24 hours, washed with PBS, and then paired with neurons cultured on CSPG+PDL or PDL matrices in artificial cerebrospinal fluid for 4 hours without direct contact. A subset of our total metabolite library was selected to assess metabolic changes in addition to the incorporation of the isotope-labeled carbon, derived from astrocytic lipid breakdown, in astrocytic and neuronal metabolic pathways (Extended Data Fig. 8).

Neurons cultured on CSPGs exhibited a marked reduction in reduced glutathione (GSH), a major intracellular antioxidant, relative to the PDL group, consistent with the findings in our 8-oxo-guanine data that CSPGs increase neuronal oxidative stress. CSPG neurons also showed increased accumulation of orotate, potentially indicating increased nucleic acid synthesis used for DNA repair or RNA maintenance (Extended Data Fig. 8a,b). Notably, dihydroorotate dehydrogenase (DHODH), the mitochondrial enzyme responsible for orotate synthesis, also plays an important role in protection against oxidative stress and ferroptosis [68], therefore indicating that this metabolic re-routing also provides antioxidant benefit. In addition, CSPG neurons displayed increased lactate and citrate, but decreased α-ketoglutarate abundance (Extended Data Fig. 8a,b).

Together, these changes indicate that CSPGs lead to a disruption in neuronal TCA cycle progression potentially through oxidative-stress mediated inhibition of aconitase [69, 70] and/or increased utilization of TCA intermediates (e.g. α-ketoglutarate) for compensatory biosynthetic functions such as glutamate replenishment or nucleotide synthesis. Within 4 hours of co-culture, neurons paired with either WT or TG2-/- astrocytes showed few significant differences (Extended Data Fig. 8b).

Astrocytes in CSPG co-cultures showed remarkably similar metabolite changes as their neuronal counterparts— increased orotate and citrate, and decreased α- ketoglutarate— and additionally exhibited reduced ketones and glutamate and increased oxaloacetate (Extended Data Fig. 8c,d). This profile suggests that within 4 hours of non-contact exposure to CSPG-conditioned neurons, astrocytes also undergo TCA cycle disruption, marked by a relatively smaller degree of citrate accumulation compared to neurons. As in the neurons, astrocytes showed comparable metabolite levels across genotype conditions within the co-culture timeframe (Extended Data Fig. 8d).

Isotope tracing in co-cultured steady state astrocytes showed that lipid-derived carbons were widely distributed throughout astrocyte metabolic pathways after 24 hours of incubation (Extended Data Fig. 8e). Exposure to CSPG-conditioned neurons stimulated an increase in astrocytic lactate labeling and a decreased flux toward α- ketoglutarate (Extended Data Fig. 8e). Isotope-labeled metabolites were transported from astrocytes to neurons during co-culture, and in PDL conditions, prominent labeling was observed in neuronal metabolites such as lactate, pyruvate, acetoacetate, α- ketoglutarate, and glutamate. Neurons on CSPGs showed increased utilization of astrocyte-derived labeled carbons for GSH synthesis and glycolytic pathways and reduced labeling of α-ketoglutarate, which further suggests impaired TCA cycle function in CSPG conditions (Extended Data Fig. 8f).

## Discussion

We demonstrate that a weight drop model of rmTBI causes persistent disruption of major white-matter tracts, reductions in spontaneous neural activity, and functional hypoconnectivity across multiple networks 28 days after injury. Structural and functional changes in these regions were associated with increased GFAP expression as well as prominent epigenetic repression and metabolic restriction in astrocytes. Additionally, we identified TG2 as a key regulator of TBI pathology and metabolic adaptability in astrocytes following injury. Whereas WT mice displayed a predominant decrease in spontaneous BOLD activity and functional connectivity after TBI, their TG2-/- counterparts showed consistent increases in both categories. This was associated with a relative preservation of large white matter tract integrity and clear reduction in reactive astrocyte GFAP expression in TG2-/- mice following injury. We hypothesize that this axonal preservation is due in large part to enhanced metabolic support by TG2-/- astrocytes following injury. Indeed, our previous studies show that astrocytic TG2 depletion facilitates neurite outgrowth in culture and functional recovery following spinal cord injury in association with upregulated metabolic pathways [42, 44]. We found that rmTBI induced significant astrocytic impairment in glutamate recycling, TCA anaplerosis, and energy production, which was attenuated in TG2-/- astrocytes.

TBIs are often characterized by shearing and rotational forces that cause multi- focal axonal injury. The white matter tracts most susceptible to injury include long coursing and commissural fibers within large scale neural networks [13, 71, 72]. The DMN, encompassing midline structures such as the posterior cingulate cortex (RSP in mice) and medial prefrontal cortex, is one such network that is frequently disrupted in TBI patients and animal models, contributing to persistent cognitive symptoms [72, 73]. Accordingly, our TBI model showed clear disruptions in major white matter tracts coincident with decreased spontaneous BOLD activity in the RSP and increased activity in medial prefrontal regions (PL and ILA) (Fig. 2a,b), the latter of which may reflect intraregional compensatory changes following disruption of long-range connections and has previously been reported [73]. It is important to note that our choice of impact site (bregma) overlies the RSP and therefore may have further exacerbated pathology in this region. The HIP, a region with extensive reciprocal connections to the DMN, also exhibited reduced spontaneous activity, accompanied by decreased functional connectivity with medial prefrontal regions (Figs. 2b, 3c). Additionally, GFAP immunoreactivity was particularly increased after TBI in DMN regions and in associated white matter connections, including the CWMB (Fig. 5g).

Among the pathological mechanisms underlying TBI, metabolic dysfunction is particularly consequential, creating a bottleneck for the energy–intensive processes required for cellular repair and network stabilization. Because the brain lacks substantial intrinsic energy reserves, neurons largely depend on astrocytes for rapidly responsive and adaptable metabolic support—both under physiological conditions and, critically, following injury [16–18]. Thus, preserving astrocyte-neuron metabolic coupling is essential for functional recovery after injury. Here, we further characterize TBI-induced metabolic dysregulation in astrocytes. WT astrocytes isolated from injured brains showed significant impairment in indicators of metabolic versatility, with pronounced downregulation in TCA- anaplerotic pathways such as lipid catabolism and glutamate transport, accompanied by decreased abundance of TCA intermediates. Impaired TCA cycle activity restricts astrocytic biosynthetic functions and limits the export of metabolic substrates required to sustain neuronal function and repair. Additionally, downregulation of EAAT2 (2.15-fold) and GLAST (1.8-fold) in injured WT astrocytes indicates a significant impairment in synaptic glutamate clearance and therefore greater vulnerability to excitotoxicity.

Conversely, TG2-/- astrocytes exhibited upregulations across metabolic pathways that contribute to TCA cycle anaplerosis and improved mitochondrial function. The metabolic upregulation observed in pyrimidine synthesis in TG2-/- astrocytes after injury is also interesting, as this is an energy-intensive pathway [74], which may further indicate an increased energy and substrate availability in this group compared to WTs. Increased pyrimidine synthesis complements the upregulation of glycogen metabolism (Fig. 7b,c) through contribution of UDP nucleotides. Additionally, we found that RNA regulatory proteins are prominently upregulated in both WT and TG2-/- astrocytes after TBI, which may be indicative of injury-related RNA dysregulation (Fig. 6g). Upregulated pyrimidine synthesis may therefore provide nucleotide substrates to support RNA maintenance. Unfortunately, few studies have examined the effect of brain injury on RNA regulation and metabolism [75]; our data suggests this should be further studied. While we identified many metabolic changes complementary with our proteomic data, our scope for assessing metabolomic changes was limited to 81 metabolites, and importantly we could not assess lipidomic profiles in our groups, which we hypothesize would be differentially regulated in TG2-/- groups.

Our *in vitro* findings reinforce TG2’s effect on astrocytic metabolic flexibility. TG2 deletion improves astrocytic support for neurons under stress, promoting neurite outgrowth [37, 42], and TG2–/– astrocytes more effectively maintain neuronal redox balance (Extended Data Fig. 6), consistent with increased lipid–utilization capacity (Extended Data Fig. 7). Under CSPG–induced oxidative stress, astrocyte–neuron co–cultures exhibited TCA cycle impairment and diversion of carbon toward orotate—a shunt that recapitulates the TBI TG2–/– profile but is absent in TBI WT astrocytes (Extended Data Fig. 7). Beyond supporting RNA and DNA maintenance, this rerouting likely enhances protection against oxidative stress through DHODH–mediated reduction of mitochondrial ubiquinone [67]. Given the short 4 hour co–culture window, longer incubations (e.g., 5 days, as used for 8–oxo–guanine and neurite–outgrowth assays) may reveal more pronounced genotype–specific metabolic differences.

Although TG2 has diverse cellular roles, accumulating evidence from our lab indicates that its epigenomic, transcriptionally repressive functions are key regulators of astrocyte reactivity after injury [28, 42, 44]. While the specific nuclear activities of TG2 that underlie transcriptional and functional changes in astrocyte injury response have not been fully defined, it likely involves a combination of catalytic and protein scaffolding functions [32, 40, 42, 43]. Among its catalytic activities, TG2 can modify histones through monoaminylation, using serotonin, dopamine, and histamine as substrates, which can facilitate or repress gene expression and alter neural activity [39, 40, 76]. Additionally, our lab has recently identified novel TG2 nuclear interactors that we hypothesize facilitate the predominant transcriptional repressive effects of TG2 in astrocytes in injury contexts— Zbtb7a, a ubiquitously expressed transcription factor, and Sin3a, an HDAC corepressor complex [37, 42]. Interestingly, by analyzing all genes with significantly reduced H3K9ac in WTs after TBI using Enrichr transcription factor enrichment, we identified Zbtb7a as one of the top significantly enriched categories (Extended Data Fig. 9a), consistent with its known repressive roles on gene transcription [77–79]. Analyzing the subset of these genes that overlapped with significant protein changes in the same group showed YY1 as the top enrichment, a transcription factor that acts as a transcriptional activator and repressor related to cell survival and injury response (Extended Data Fig. 9b) [80]. Overlap with protein changes in the TG2-/- astrocytes after TBI showed Zbtb7a as the most significant transcription factor enrichment (Extended Data Fig. 9c), and this enrichment largely represented upregulated proteins, supporting our hypothesis that TG2 depletion alleviates Zbtb7a-associated transcriptional repression in injury contexts. We propose that TG2 may facilitate transcriptional repression in astrocytes by scaffolding Zbtb7a and Sin3a, potentiating recruitment of histone deacetylase to Zbtb7a-targeted metabolic genes. Further studies are needed to elucidate this mechanism and the potential contributions from TG2-catalyzed histone modifications.

While constitutive TG2 deletion led to striking pathological differences associated with astrocyte metabolism following rmTBI, the use of male TG2–/– mice raises important considerations to address related to sex, potential contributions from other cell types, and development, particularly given the observed baseline differences between WT and TG2–/– mice. Therefore, the mechanistic working model we have developed in this study serves as a foundation for future studies using astrocyte–specific conditional TG2 deletion in male and female mice to more precisely define the contribution of astrocytic TG2 to TBI outcomes. These studies also highlight TG2 as a promising therapeutic target; indeed, we previously showed that TG2 inhibition in spinal cord injury and in *in vitro* neurite outgrowth phenocopies the effects of astrocytic TG2 deletion [44]. Future work will therefore determine whether pharmacological TG2 inhibition can reproduce the protective metabolic changes observed in TG2–/– astrocytes to mitigate persistent TBI pathology.

## Methods and Materials

### Animals

All animal procedures were reviewed and approved by the University Committee on Animal Resources of the University of Rochester. Animals were maintained on a 12 h light/dark cycle with access to food and water ad libitum. WT C57BL/6N mice were originally purchased from Charles River Laboratories. Our TG2−/− mice on a C57Bl/6N background were described previously and have been continuously bred in house [26].

### Repetitive mTBI (rmTBI) weight drop model

Male wild type and TG2-/- mice, 10-12 weeks old, were used for rmTBI. A modified weight drop model was used deliver mTBIs as previously characterized [47]. In brief, mice were anesthetized with 3% isoflurane, placed on a sterile pad over a foam platform, secured with two elastic straps, and a 444 g weight was dropped, hitting the impactor, which is installed with a Teflon tip that diffuses the impact force over the skull. Anatomical bregma was targeted by positioning the center of the impactor at the midline point at the anterior base of the ears. The impactor was driven downward into the skull at 3.1m/s velocity, delivering an approximate force/area of 3.3 N/mm^2^. Following the impact, a spring and solenoid piston work in concert to “catch” the weight, allowing the animal’s skull to rapidly return to its original position, producing a whiplash effect with combined linear and angular accelerations, which were quantified from both an accelerometer fixed to the weight sled and high-speed videos taken of every impact [47]. Immediately after the impact the mice were placed in separate cages in supine position until recovery. The time to right themselves from the supine position was recorded.

#### Quantification of angular acceleration

High-speed videos from 24 representative TBI impacts were analyzed using Kinovea motion tracking software [49]. Frame-by-frame tracking was used for accurate capture of angular displacement across time. Digital markers for frame-by-frame tracking were placed at the nose tip, right ear midpoint, and bottom of a fixed ruler throughout all videos. Angular acceleration graphs were then generated in Kinovea.

### Immunohistochemistry

In preparation for immunohistochemistry, animals (n= 4 per group) were perfused with ice-cold PBS, and their brains were immediately extracted and submerged in 4% paraformaldehyde for 4 hours. Brains were then placed in a 10/20/30 percent sucrose gradient with sequential overnight incubations. Brains were flash frozen in dry ice and cryosectioned into 25 µm sections on a freezing-stage sliding microtome. Sections were stored in cryoprotectant at −20 °C until further processing. For GFAP and NeuN staining, sections were mounted onto Superfrost Plus micro slides (VWR, 48311-703) and received three washes with PBS + 0.05% Tween 20 (PBS-T), followed by incubation with 0.25% Triton-X in PBS for 10 minutes. Slides were then submerged in citrate buffer (10mM sodium citrate, pH 6) and placed in a steamer for 40 minutes for heat-induced antigen retrieval. The M.O.M. (Mouse on Mouse) immunodetection kit (Vector Laboratories, BMK-2202) protocol was used with GFAP (GA5 clone) anti-mouse primary antibody (Cell Signaling, 3670; 1:1000). For NeuN, slides were incubated in 5% BSA in PBS-T blocking buffer for one hour followed by overnight incubation with NeuN anti-rabbit primary antibody (Cell Signaling, 12943S; 1:250). On the following day, GFAP and NeuN samples were washed and incubated with secondary antibodies in blocking buffer for one hour— streptavidin 488 (Biolegend, 405235; 1:1000) for GFAP M.O.M. samples and anti-rabbit Alexa Fluor 594 (Invitrogen, A21207; 1:1000) for NeuN samples— and samples were prepared for imaging using glycerol mounting medium (EMS, 17989-40).

#### Image acquisition and analysis

Confocal images were acquired with an Olympus FV1000MP microscope using a 40× oil objective (UApo; NA= 1.35). Image stacks were taken with a 1 µm Z-step size and 1600 × 1600-pixel resolution. All parameters (gain and intensity) stayed consistent across samples. Images were analyzed using Fiji Image J [81]. For GFAP data, areas within select brain regions were outlined, images were thresholded using Isodata automatic threshold plugin, and percent area of thresholded signal was measured. For NeuN data, select brain regions were outlined and both mean fluorescent intensity and percent thresholded (Isodata) area were measured.

### Cell Culture

Primary astrocytes were cultured between post-natal days 0 to 1 (P0–P1) from either WT or TG2-/- C57BL/6N mouse pups as previously described [26]. In brief, mouse pups were rapidly decapitated, their brains were collected and dissected, meninges removed, and cortical hemispheres were collected. Cortical tissues were triturated and dissociated cells were plated onto culture dishes in MEM supplemented with 10% FBS (Atlas Biologicals, Fort Collins, CO, USA, F-0500-DR), 33 mM glucose, 1 mM sodium pyruvate (Gibco, Grand Island, NY, USA, 11360-070), and 0.2% Primocin (Invivogen, San Diego, CA, USA, ant-pm-05) (glial MEM). Twenty-four hours after plating, the dishes were shaken vigorously and rinsed to remove debris and non–astrocytic cell types. Astrocytes were maintained at 37 °C and 5% CO2 for 7–8 days, then frozen in glial MEM with 10% DMSO and stored in liquid nitrogen. For experimental use, astrocytes were rapidly thawed and cultured in glial MEM, and cultures with 2-3 passages and no more than 90% confluency were used for final data acquisition.

Primary cortical neurons were prepared from Sprague-Dawley rat embryos at embryonic day 18 (E18) and cultured as previously described with minor modifications [82]. Coverslips were prepared by incubating poly–D–lysine (Sigma, P6407) diluted in PBS to 20 µg/mL for 4 h. After rinsing, coverslips were either stored in PBS or further coated overnight with chondroitin sulfate proteoglycans (CSPGs; Millipore, CC117; 2.5 µg/mL in PBS). All coverslips were rinsed with PBS prior to cell plating. To prepare the neurons, pregnant rats were euthanized using CO2, followed by rapid decapitation in accordance with NIH Animal Research Advisory Committee guidelines. Embryos were removed and decapitated, and cerebral cortices were dissected and cleared of meninges. Cerebral cortices were then digested in 0.05% trypsin-EDTA (Corning, 25-053-Cl) for 15–20 min at 37 °C, then gently triturated. Dissociated neurons were plated at 12,000 cells/cm^2^ in MEM (Gibco, 42360032) supplemented with 5% FBS, 20 mM glucose, and 0.2% Primocin. Four to five hours after plating, the medium was replaced with Neurobasal medium (Gibco, 21103-049) containing 2% B27 (Gibco, 17504-044), 0.5 mM Glutamax (Gibco, 35050-061), and 0.2% Primocin (neuron growth medium). Neurons were maintained at 37 °C and 5% CO2.

### Immunocytochemistry

Primary astrocyte and neuron cultures were washed in PBS and fixed in 4% paraformaldehyde/4% sucrose for 5 min, followed by two PBS washes and permeabilization with 0.25% Triton–X for 10 min. For 8–oxo–guanine detection, cells were incubated in 1.5 M HCl for 30 min to relax DNA and enable antigen retrieval, neutralized in 0.1 M NaOH for 5 min, and washed three times before treatment with RNase (Thermo Fisher EN0531) at 37 °C to remove oxidized RNA; negative–control wells were additionally incubated with DNase (Sigma D5025). After two further washes, cells were blocked in 5% BSA for 1 h and incubated overnight with anti–8–oxo–G mouse antibody (Santa Cruz sc–130914; 1:400), followed by goat anti–mouse Alexa Fluor 594 (Invitrogen A–11032; 1:1000) for 1 h. For mitochondrial morphology assays, astrocytes were blocked in 5% BSA and immunolabeled with anti–TOM20 rabbit antibody (Proteintech 11802–1–AP; 1:500) and goat anti–rabbit Alexa Fluor 488 (Jackson Lab 111–545–045; 1:1000).

### Cortical astrocyte isolation

Cortical astrocytes were isolated from sham and TBI mice 1 month after the rmTBI protocol, following MRI. Magnetic-activated cell sorting (MACS) was used with anti- ASCA2-micro magnetic beads (Miltenyi, 130-097-678) to select for astrocytes, modified from a validated protocol. Briefly, mice were anesthetized by isoflurane and rapidly decapitated. Brains were extracted and cortical tissue was dissected out from the rest of the brain and minced to roughly 1 cubic millimeter pieces. Tissue dissociation was performed using papain (Worthington Biochemical, LK003178) diluted in homemade artificial cerebral spinal fluid (aCSF) (125 mM NaCl, 5 mM KCl, 2 mM MgSO_4_, 24 mM NaHCO_3_, 1.25 mM NaH_2_PO_4_, 2 mM CaCl_2_, 10 mM glucose, pH 7.4), plus DNase I (Sigma, D5025), followed by gentle trituration, centrifugation, supernatant removal, and resuspension with aCSF buffer containing ovomucoid inhibitor (Worthington Biochemical, 50-592-496). Tissue was further triturated, centrifuged, supernatant was removed, and the pellet was resuspended in 5% BSA in PBS. The suspensions were filtered through 40 µm cell strainers followed by MACS protocol for astrocyte isolation using LS columns (Miltenyi, 130-042-401). Astrocytes were eluted from MACS LS columns in aCSF, pelleted, and frozen at −80°C for CUT&RUN, proteomic, and metabolomic analyses.

#### CUT&RUN

Each isolated astrocyte sample (n= 2 per group) was resuspended in 500 μl of nuclear extract (NE) buffer (20mM HEPES-KOH, pH 7.9, 10mM KCl, 0.5mM Spermidine, 0.1% TritonX-100, 20% glycerol, freshly added protease inhibitors (Halt Protease and Phosphatase Inhibitor Cocktail, 78446)) [83] for nuclear isolation in DNA LoBind 1.5 ml microfuge tubes (Eppendorf, 13-698-791). Sodium butyrate (10mM) was added to buffers including protease inhibitors to inhibit histone deacetylase activity during processing. Nuclei were pelleted at 1,100g for 5 min at 4 °C in a swinging-bucket rotor and supernatant discarded. Nuclei were washed again in 500 μl NE buffer and counted using a hemocytometer. In total, 100,000 nuclei were used per biological replicate.

BioMag Plus Concanavalin A beads (Polysciences, 86057) were prepared (15 μl bead slurry per reaction) by washing three times in 500 μl binding buffer (20 mM HEPES-KOH, pH 7.9, 10 mM KCl, 1 mM CaCl_2_, 1 mM MnCl_2_) and resuspending to 500 μl binding buffer [83–85]. For all CUT&RUN reactions, including IgG negative controls, 15 μl of the resuspension was aliquoted into 1.5 ml DNA LoBind tubes (Eppendorf, 022431021) and 500 μl NE buffer was added to each tube. 100,000 isolated nuclei from cells were added to each tube, and rotated end over end at room temperature for 10 min. The bead-bound nuclei were washed with 1 ml wash buffer (WB: 20 mM HEPES, pH 7.5, 150 mM NaCl, 0.1% Triton X-100, 0.1% Tween-20, 0.5 mM spermidine, 0.1% BSA, freshly added protease inhibitors, 10mM sodium butyrate) three times. All washes were done with minimal pipetting, and beads were mixed by inversion and light flicking of the tube. Beads were resuspended in 100 μl of antibody buffer (1 ml wash buffer with 2 mM EDTA) and mixed by flicking. Two microliters of H3K9ac antibody (Millipore Sigma, 07-352) or CUTANA IgG negative control (EpiCypher, 13-0042) (1:50) were added to their corresponding tubes and mixed by flicking. Bead-bound nuclei were incubated on a mixer (tubes on their side at ∼20-degree upward angle) with primary antibodies overnight at 4 °C. Nuclei were washed twice the next day with 1 ml of cold WB. Nuclei were resuspended in 50 μl of cold wash buffer by flicking, and 2.5 μl of pAG-MNase (Epicypher, 15-1016) was added and mixed by flicking and incubated for 1 h at 4 °C on the same mixer. Nuclei were then washed three times with 1 ml ice-cold wash buffer, followed by one wash in 1 ml low-salt rinse buffer (20 mM HEPES, pH 7.5, 0.5 mM spermidine, 0.1% Tween-20 and 0.1% Triton X-100). Nuclei were resuspended in ice-cold calcium incubation buffer (3.5 mM HEPES, pH 7.5, 10 mM CaCl_2_, 0.1% Tween-20, 0.1% Triton X- 100) and immediately placed into an ice-cold metal block on ice to maintain the temperature. The samples were incubated for 30 min, and then 100 μl of 2× stop buffer (340 mM NaCl, 20 mM EDTA, 5 mM EGTA, 0.1% Tween-20, 0.1% Triton X-100, 25 μg/ml

RNase A (Thermo Fisher Scientific, EN0531)) and 0.1 ng per 100 μl of E. coli spike-in DNA (Epicypher, 18-1401) was added, and the beads were mixed by flicking. Nuclei were incubated at 37 °C for 15 min with no shaking to allow for release of chromatin and digestion of RNA. Beads were placed onto a magnet, and the supernatant (200 μl) was collected. The supernatant was treated with proteinase K for 1 hour at 50°C to degrade all proteins. DNA was isolated using the Zymo ChIP DNA Clean & Concentrator kit (D5205) and eluted in 30 μl and frozen at −20 °C for library preparation [40].

#### CUT&RUN-seq library preparation and sequencing

Library preparation was performed using the NEBnext Ultra II DNA library kit (E7645L) with multiplexed adapters with minor modifications. CUT&RUN DNA underwent end repair and adapter ligation according to the manufacturer’s protocol (1:15 adapter dilution was used). DNA was amplified using 16 PCR cycles with 10 s of extension time per cycle. Libraries were quantified using the Qubit fluorometer (Thermo Fisher Scientific) DNA high sensitivity kit, and the library size distribution was checked using the Tapestation DNA High Sensitivity ScreenTape (Agilent). Libraries were pooled at an equimolar concentration and sequenced on the Illumina NovaSeq X+ sequencer by the NYU Genome Technology Center (paired-end, 100 cycles).

#### CUT&RUN–seq data analysis

Raw sequencing files were demultiplexed using bcl2fastq2 (Illumina, v.2.20). Between 20 and 100 million total reads were achieved for each replicate (average, 42.6 million). The samples were aligned to the mm10 genome using bowtie2 (v.2.5.0) [86], with the following parameters: --local --very-sensitive-local --phred33 -I 10 -X 700 --dovetail --no-unal --no-mixed --no-discordant [84]. Low-quality reads were filtered out using Samtools (v.1.9) with a cut-off MAPQ score of 30, and only unique reads were retained for further processing [87]. Unique read files for each replicate/timepoint/antibody were merged and used for peak calling using MACS2 (v.3.0.0a6) with the callpeak function and the options -f BAMPE -q 0.05 --broad --broad-cutoff .05, using the corresponding IgG sample as the - c. For visualization, each sample was normalized by scaling the samples based on the E. coli spike-in DNA. Each sample was aligned to the E. coli genome (MG1655), and the uniquely aligned reads were counted. The number of E. coli reads for each replicate between timepoints was compared, with the sample with the lowest number of E. coli reads set at a scaling factor of 1×. The other samples were scaled down by a scaling factor that was computed by dividing the lowest number of E. coli reads by the sample number of E. coli reads. Genome coverage tracks (bigwig files) were produced using the deepTools (v.3.5.1) bamCoverage function with the options --binSize 10 --smoothLength 30 --normalizeUsing None --scaleFactor # (derived from E. coli spike in) and using an ENCODE mm10 blacklist file (https://doi.org/10.1038/s41598-019-45839-z, v2) to discard regions with consistently non-specific signal [88]. Deeptools multibigwigSummary was run genome wide using a sliding window (window size 500 basepairs, offset by -450 basepairs), and the significant regions identified by comparison to the IgG controls (using a minimum cutoff of 20 reads for H3K9ac and 10-fold increase over IgG). Overlapping and neighboring regions were merged using Bedtools (merge -d 501). Then, the average number of H3K9ac reads per peak was calculated using multibigwigSummary. Overlap of various peaks and TSSs was achieved using bedtools intersect (v.2.31). Heatmaps were made using Deeptools (v.3.5.5) computeMatrix and plotHeatmap in reference-point mode, centered over TSSs or peak centres, using binSize 10 and --sortUsing mean, sorted in descending order. TSSs were downloaded from the UCSC (mm10) table browser using the canonically annotated transcript for each gene. The resulting matrix files were analyzed in R using dplyr (v.1.1.4), calculating the ratio between sham and injured animals in both genotypes. Genes with ≥ 1.5 and < 1.5-fold change comparing sham to TBI samples were plotted as heatmaps using the same deeptools parameters as above.

### Proteomic Analysis

Astrocyte proteomic samples (n= 6 per group over 3 separate batches) were processed by the University of Rochester Mass Spectrometry Resource Lab and analyzed using data independent acquisition in an Orbitrap Fusion Lumos Tribrid quantitative mass spectrometer (Thermo Fisher Scientific), as previously described [42].

#### Protein Differential Expression Analysis

Differentially expressed proteins were identified using pairwise *t*-tests followed by Benjamini–Hochberg (BH) FDR correction. Data were sorted by log2 fold change and adjusted *p* value. Log2 fold change values between 0.5 and −0.5 and adjusted *p* values > 0.05 were excluded from further analysis. Filtered data across separate batches were collated and duplicate gene IDs were removed. Figures showing overall protein changes were generated in Excel.

### Gene Ontology Analyses

Sorted differentially expressed gene symbol lists from CUT&RUN and proteomic data were analyzed using DAVID GO Biological Process and KEGG pathway enrichment [89]. The enrichment data was extracted for generating dot plots using ggplot2 package in R 4.5.3. Gene lists were also analyzed by Enrichr for transcription factor and cell type enrichments [90–92].

### Metabolomic Analysis

#### Isolated cortical astrocytes

Frozen astrocyte cell pellets in 2 ml tubes were resuspended in 750 µL of ice-cold 80% methanol and homogenized in Dounce glass homogenizers for metabolites extraction. Samples were centrifuged at 14,000 rpm for 5 minutes and supernatants were transferred to new 2 ml tubes. Extracts were dried under N2 stream and resuspended in 150 μL of 50% methanol. Pool samples were created and analyzed alongside original samples for HPLC run quality control. All 4 experimental groups were analyzed together in 4 batches over ∼6 months, with two biological replicates per group in each batch. Replicates were arranged in staggered order throughout the run, for a total of 10 samples analyzed per run (8 samples and 2 pools at start and end). From resuspended extracts, 15 µL was injected onto a Synergi Fusion-RP column (Phenomenex) for reverse phase LC separation in a Prominence 20A HPLC system (Shimadzu), and a 3% to 100% methanol ramp was performed over 50 min. HPLC effluent was subject to electrospray ionization in negative ion mode using a HESI-II ion source. Samples were analyzed by single reaction monitoring on a Thermo Quantum triple-quadrupole mass spectrometer. Metabolites were identified against a library of validated standards based on retention time, intact mass, collision energy, and fragment masses, allowing for detection of ∼120 common metabolites [93]. Data was analyzed using Thermo XCalibur 4.0 software, and 81 metabolites were reliably detected across samples.

For all batches, metabolite relative abundances were normalized by the sum of all abundances within each sample. Batches were then normalized to each other on a per-metabolite basis using average (relative) metabolite abundance across all groups within a batch. Batches were then aggregated and outliers were identified using 99.99% confidence intervals for each group. Outliers and missing values (in total 106 of 2592 possible data points) were imputed on a per-metabolite basis from medians of remaining values within each group [94]. Data was analyzed with the Real Statistics Resource Pack software (Release 9.7.6) in Excel, using pairwise t-tests, and p values <0.05 were considered significant.

#### Isotope Tracing

Astrocytes cultured on 30mm coverslips (Bioptechs, 30-1313-0319) were incubated with 50 µM U-^13^C palmitate (Cambridge Isotope Laboratories, CLM-409-0.01) in substrate limited media (Gibco MEM + Glutamax, 41090036, supplemented with 10% FBS) for 24 hours. Astrocyte cultures were then paired with neuron cultures by laying the astrocyte coverslip over paraffin pedestals embedded on the neuron coverslip (adapted from Ioannou et al.). Astrocytes and neurons were paired for 4 hours, and then the coverslips were separated into 6 well culture plates on ice, quickly washed in ice-cold PBS, and immersed in 750 µL of ice-cold methanol for metabolite extraction as described above. Data was processed as above, however outliers and missing values were not imputed in this dataset.

### MRI experimental design

Animal preparation and MRI data acquisition: Animals (n= 6 per group) were briefly exposed to isoflurane for placement into the restrainer, and attachment of physiological monitoring devices. To avoid the inhibitory effects of isoflurane on the blood oxygen level-dependent (BOLD) signal, resting state functional MRI (rsfMRI) was acquired under dexmedetomidine (MD - Dechra Pharmaceuticals, USA) sedation [95, 96]. Moderate MD sedation was initially induced by a subcutaneous bolus injection (0.3 mg MD/kg body weight in 100 μL 0.9% NaCl solution); 15 min later, the animals received a continuous infusion of MD through an MRI-compatible catheter (0.6 mg/kg body weight in 200 μL/h) subcutaneously inserted at the mouse abdominal level. rsfMRI data were acquired using T2*-weighted single-shot gradient-echo echo-planar imaging (GE-EPI) sequence with a 9.4T small-bore animal scanner (Biospec 94/30, Bruker, Germany) and a cryogenically cooled mouse head adapted CryoCoil. Four hundred volumes were recorded in an interlaced manner for each ∼10min run using the following parameters: TE/TR=14/1500 ms, 19 axial slices of 0.5 mm thickness, with an FOV of 19.2×12 mm^2^ and an acquisition matrix of 128×80, resulting in a planar resolution of 150×150 μm^2^.

Following rsfMRI, MD infusion was discontinued and morphological images were acquired using a multi-slice T2-weighted Turbo-RARE sequence [97] (TE/TR = 8 ms/2500 ms, two averages, RARE factor of 8). The entire mouse brain was imaged using 25 slices of 0.5 mm thickness with a FOV of 20 × 20 mm^2^, and acquisition matrix of 256 × 256, resulting in a planar resolution of 78 × 78 µm^2^.

Finally, diffusion MRI (dMRI) data was acquired under isoflurane (∼1.5 vol%) using respiration triggering to avoid motion-related artifacts. Multi-shell dMRI data were acquired using a four-shot multi-slice diffusion tensor imaging-EPI (DTI-EPI) sequence [95, 96] with b factors of 1000 s/mm^2^ and 2000 s/mm^2^ in 45 noncollinear diffusion gradient directions per b-value. 25 axial slices of 0.40 mm thickness were acquired at a spatial resolution of 100 × 100 μm^2^ with an FOV of 12 × 8 mm^2^ and an acquisition matrix of 120 × 80, and the following diffusion parameters: TE/TR = 25 ms/3500 ms, diffusion gradient separation (Δ) = 10 ms, diffusion gradient duration (δ) = 4.0 ms. Physiological conditions (temperature and respiration) were continuously monitored during the imaging session using a small animal monitoring system (SAI, Inc., Stony Brook, NY).

### rsfMRI data pre-processing

rsfMRI data were reconstructed and preprocessed using MATLAB R2025a (www.mathworks.com) scripts developed in-house [98, 99]. In brief, the following steps were applied for each animal: Despiking: The first ten volumes of each raw rsfMRI dataset were removed to ensure the steady state of magnetization. Motion outliers were then identified by computing framewise displacement (FD) as described in [98, 100]. Volumes with FD > 0.2 mm, along with their adjacent volumes, were removed.

Image coregistration: rsfMRI and dMRI datasets were coregistered to the template using the image coregistration pipeline described previously [51, 95, 101, 102]. First, rsfMRI data were aligned to the respective T2-weighted image volume using a normalized mutual information approach, 4th degree B-Spline interpolation, and a six-parameter rigid body transformation (three translations and three rotations). The rsfMRI and corresponding T2 data were then transformed into a template space (150 µm³ isotropic resolution) aligned with the mouse atlas [51] using automated image registration (AIR) implemented in DiffeoMap (www.mristudio.org).

Motion correction: Motion artifacts were corrected by realigning rsfMRI volumes to the first volume using a least-square approach with a six-parameter rigid body spatial transformation, as implemented in statistical parametric mapping (SPM) [95, 98].

Nuisance regression: White matter and cerebral spinal fluid masks from the atlas were mapped onto each subject’s rsfMRI data, and nuisance signals from these compartments, together with motion parameters, were regressed out voxel-wise from the BOLD time series [98].

Smoothing and filtering: The resulting rsfMRI data were spatially smoothed (Gaussian kernel, FWHM = 1 mm) and temporally filtered by a 0.01–0.1 Hz bandpass filter [95, 98].

### dMRI data pre-processing

dMRI data were pre-processed as described previously [51, 99, 101]. Briefly, non-brain tissue signals were manually discarded in AMIRA (ThermoFisher Scientific, https://www.thermofisher.com) and ghosting artifacts, likely caused by frequency drifts, were corrected using rigid alignment in DTIStudio [103].

### Anatomical labeling of MRI data

Both rsfMRI and dMRI datasets were coregistered to the high-resolution dMRI-based mouse brain atlas [51] using rigid transformation. Detailed structural labels containing 42 brain regions were transferred to each subject, as described previously [51, 101]. These subject-specific regions were used to compute functional and structural metrics for each mouse.

### MRI data post-processing

#### ALFF

ALFF was used to quantify spontaneous neural activity at rest by measuring BOLD fluctuations within the 0.01-0.1 Hz range [50]. Voxel-wise time series were transformed into the frequency domain using a fast Fourier transform, and ALFF was measured by computing the square root of the power spectrum and averaging throughout the bandpass (0.01–0.1 Hz) at each voxel. To standardize across subjects, voxel-wise ALFF values were normalized by the global ALFF value. Group-level ALFF maps were then generated [99, 104], and atlas-defined regions were used to assess genotype- and injury-dependent effects on regional ALFF.

#### rsFC

Twenty-one bilateral atlas-defined brain regions (nodes) spanning the whole brain were used to compute node-to-node rsFC [51]. The time course of each node was obtained by averaging the time courses of all voxels within the node. Node-to-node connectivity was calculated using Pearson correlation between regional time courses and converted to z-scores using Fisher’s r-to-z transformation.

#### FA

We computed FA to examine the degree of overall directionality of water diffusion, which reflects the white matter integrity in the brain (Figure 4a,b). Six diffusion tensor elements were determined by log-linear fitting via DTIStudio [103]. Next, three eigenvalues (λ1, λ2, λ3) and corresponding eigenvectors (v1, v2, v3) were computed by diagonalizing the tensor, and FA was then calculated from the eigenvalues [105, 106] as follows: √3/2 × [√ ((λ1 − λ)2 + (λ2 − λ)2 + (λ3 − λ)2) / √ (λ12 + λ22 + λ32)], where λn = the eigenvalues describing the diffusion tensor, and λ is the mean diffusivity ((λ1 + λ2 + λ3)/3). Higher FA values indicate stronger white matter integrity and vice versa.

### Statistical analyses

GraphPad Prism version 10.2.2 for Windows (GraphPad Software, La Jolla, California, USA) was used to report raw data and perform statistical analyses. Immunohistochemistry and *in vitro* 8-oxo-guanine and mitochondrial data were examined using two-way ANOVA with Šidák’s multiple comparison test with genotype and rmTBI as fixed factors. CUT&RUN data were examined using Scheirer–Ray–Hare test with post-hoc Wilcoxon-ranked sum test. Metabolomic data was examined using multiple pairwise t-tests with significance set at P< 0.05.

#### MRI statistical analysis

Statistical analyses were performed and visualized using MATLAB R2025a and GraphPad Prism. Data were examined using a two-way ANOVA with genotype and rmTBI as fixed factors to assess the effects of TG2, rmTBI, and their interaction on FA, ALFF, and rsFC, followed by Tukey’s HSD or Holm-Šidák post-hoc analysis. Our primary dependent variables (i.e., regional ALFF, regional FA, and rsFC) were examined for outliers defined as >2 standard deviation above or below the mean. We did not identify any outliers.

## Acknowledgements

The authors would like to thank Joseph Irvin and Peter Girardi for assistance with cell culture and analyses of fluorescent images, and Dr. David, Yule for use of his confocal microscope. The fee for service core facilities that were used were CALMN (RRID:SCR_023177) and the Proteomics Core (RRID:SCR_023106). These studies were supported by 2025-2026 CABIN Pilot Grant (TD, GVWJ), R01DC021213 (JPM), R21NS119673 (GVWJ), F32MH140478 (B.H.W.), R01MH116900 (I.M.), Del Monte Institute Pilot Project Program grant (GVWJ, TA, JPM), Joan Wright Goodman 2025-2026 Dissertation Fellowship (TD).

## Extended Data Figures

**Extended Data Figure 1.**
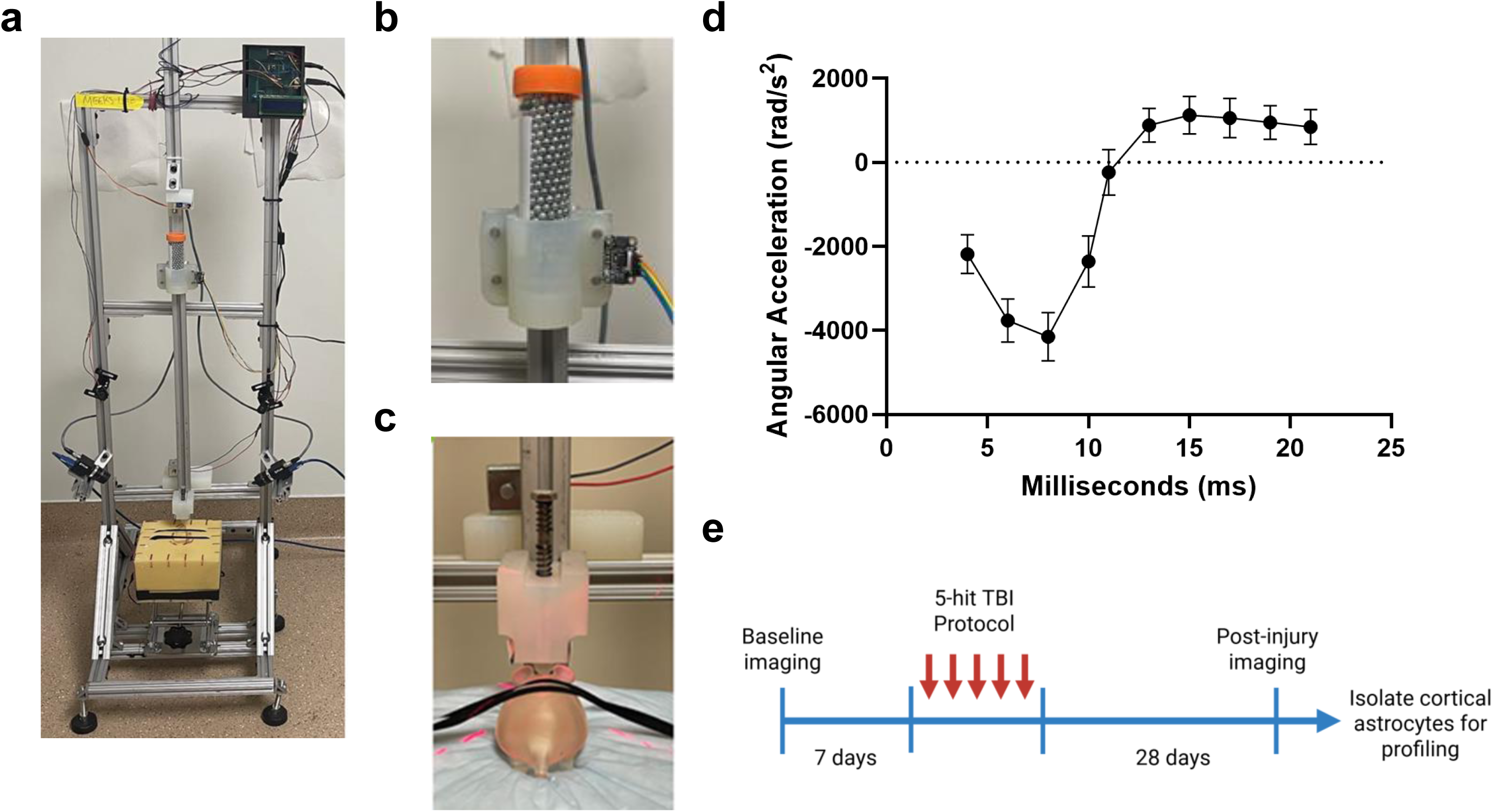
rmTBI model. **a)** Modified weight drop device showing high speed cameras and foam platform for mouse. **b)** The weight sled contains a conical tube filled with 444 g of steel airgun shot, and the sled is fixed with an accelerometer. **c)** Close-up view of the solenoid, positioning lasers, and the piston, positioned midline over the head of the 3D mouse model at the anterior base of the ears. **d)** Angular acceleration trace of the mouse head following impact, shown as mean ± s.e.m. across 27 replicates of representative TBIs, aligned to the moment of impact. **e)** Timeline of 5-hit mTBI protocol with baseline and post-injury MRI, after which the mice were sacrificed for cortical astrocyte isolation. (Images of TBI rig from Crum et al., 2026)[47].

**Extended Data Figure 2.**
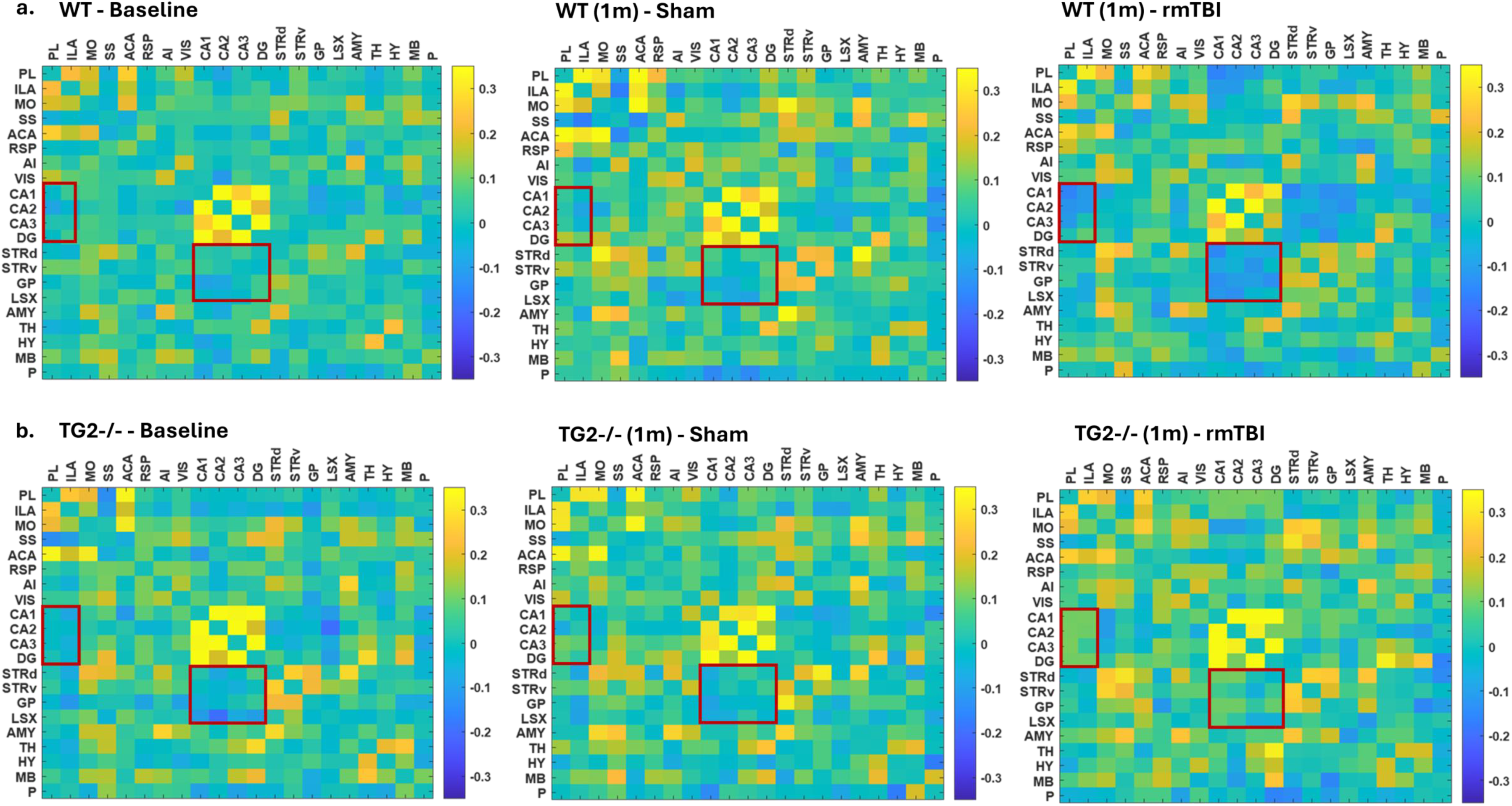
Functional connectivity matrices. Group average whole brain functional connectivity matrices of the **a)** WT and **b)** TG2-/- groups at baseline and following rmTBI or sham respectively derived via partial correlation analysis. Each cell of the matrix represents the functional connectivity between seed (row) and target (column) regions. Hot and cold colors represent enhanced or suppressed functional connectivity. (n= 6 mice per group).

**Extended Data Figure 3.**
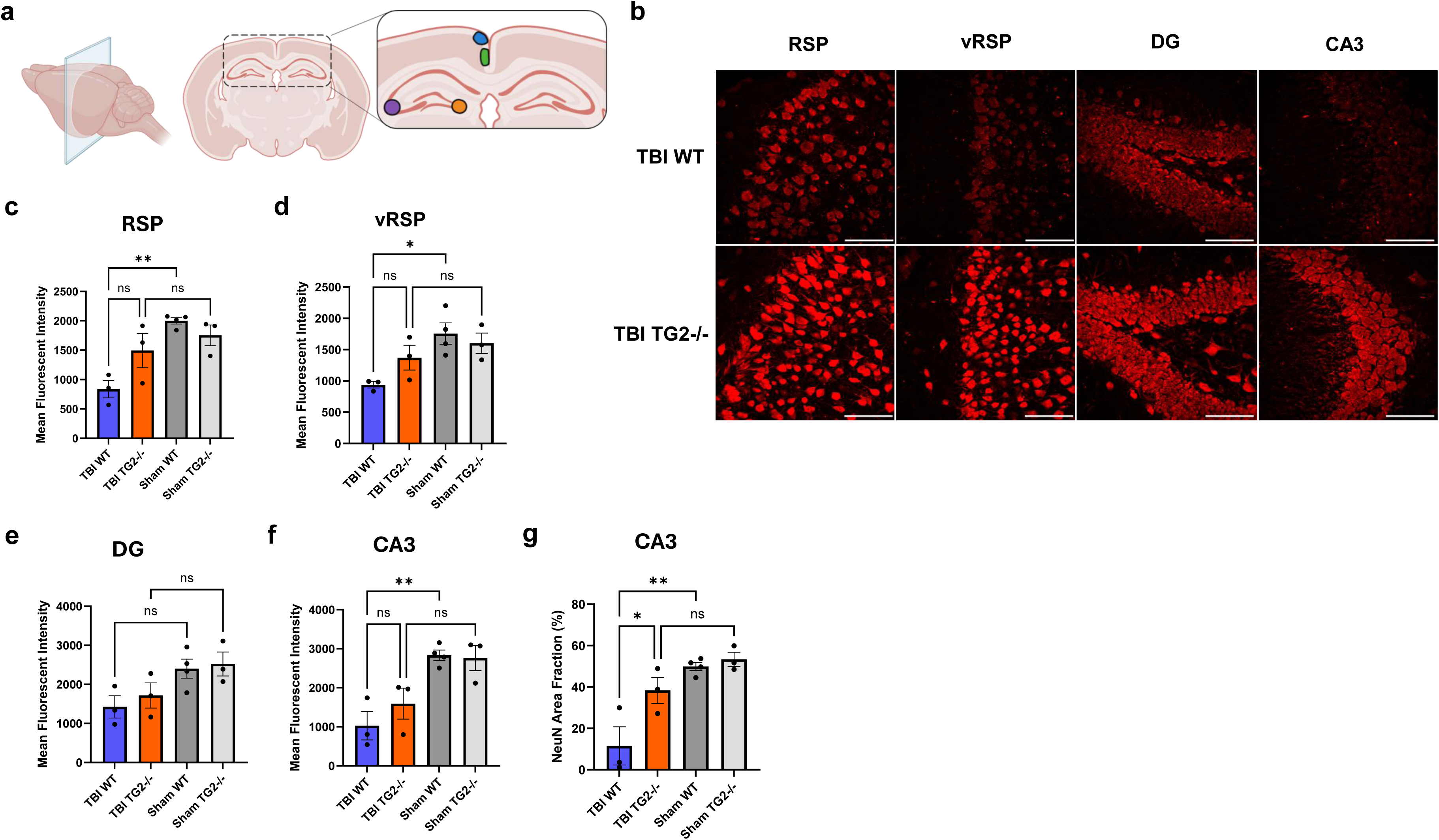
Injured WT but not TG2-/- mice show significantly decreased NeuN immunoreactivity across TBI-vulnerable brain regions. **a)** Regions within coronal sections selected for quantification of NeuN immunofluorescence (colored areas); blue– RSP, green– vRSP, purple– CA3, orange– DG. **b)** Representative 40x images (scale bars = 70 μm) and **c-g)** quantification of NeuN immunoreactivity in RSP, vRSP, CA3, and dentate gyrus (DG) across groups. Data in bar graphs represent mean ± s.e.m. (n= 3-4 mice per group, two-way ANOVA with Šidák’s multiple comparison test, α ≤ 0.05. *P < 0.05, **P < 0.01, ***P < 0.001, ns, not significant). Diagram in **(a)** created in BioRender.com.

**Extended Data Figure 4.**
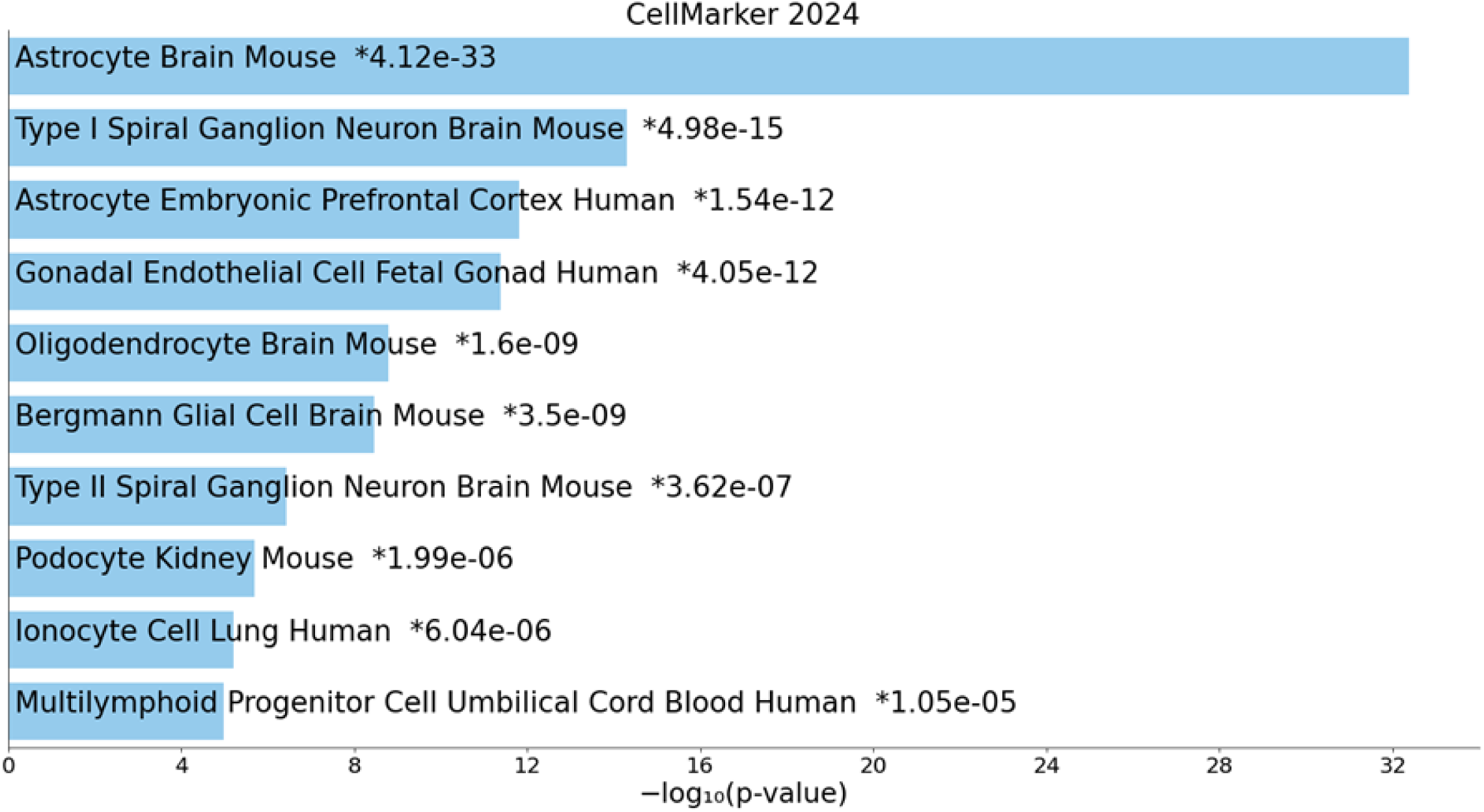
Cell type enrichment analysis validates astrocyte selectivity of ASCA2+ cortical cell isolation. Genes with ≥1.5-fold-change reduction in H3K9ac in TBI WT group were used for Enrichr analysis using the CellMarker 2.0 database. Categories are ranked by p value. Asterisks represent categories with Benjamini Hochberg corrected p value <0.05.

**Extended Data Figure 5.**
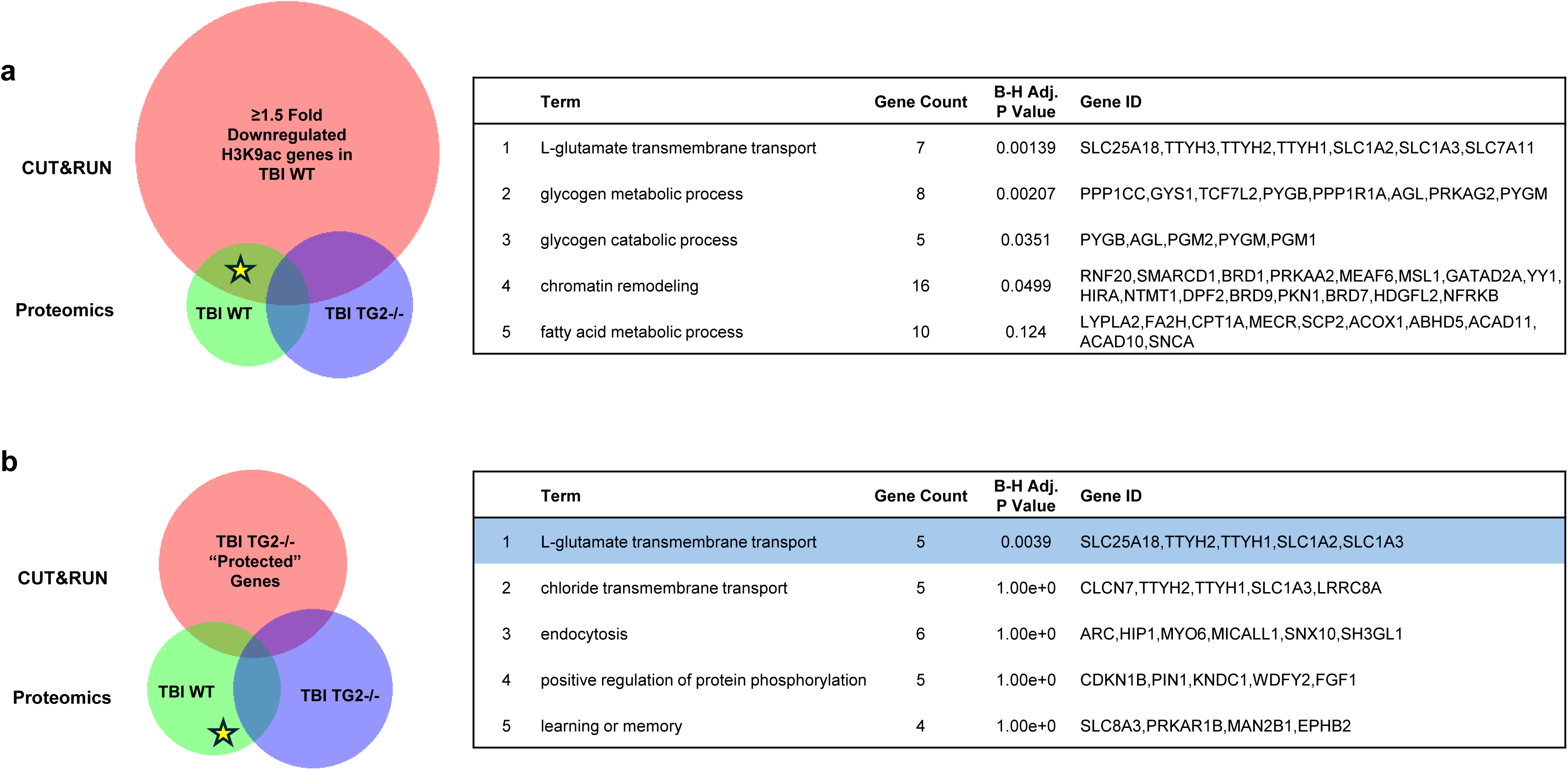
Prominent metabolic pathway protein downregulations are associated with removal of H3K9ac marks at corresponding gene loci. **a)** Venn diagram shows genes with ≥1.5-fold-change reduction in H3K9ac in TBI WT group (5607 genes) that overlapped with differentially regulated proteins in TBI WT (930 proteins) and TBI TG2-/- (1320 proteins). Enrichment analysis using DAVID GO Biological Process of TBI WT genes that were shared in both datasets (star, 381 genes) revealed that metabolic pathways which were strongly downregulated in the proteomic data, glutamate transport and glycogen metabolism (Fig. 6g), are also associated with transcriptional repression by H3K9ac reduction. **b)** The same analysis was used for genes in the TBI TG2-/- group which did not show reduction in H3K9ac (“protected”) (1844 genes) and TBI WT differentially regulated proteins (star, 131 genes). This identified glutamate transport as a pathway which may avoid transcriptional repression by H3K9ac reduction in TG2-/- groups after injury. Venn diagrams generated using BioVenn.

**Extended Data Figure 6.**
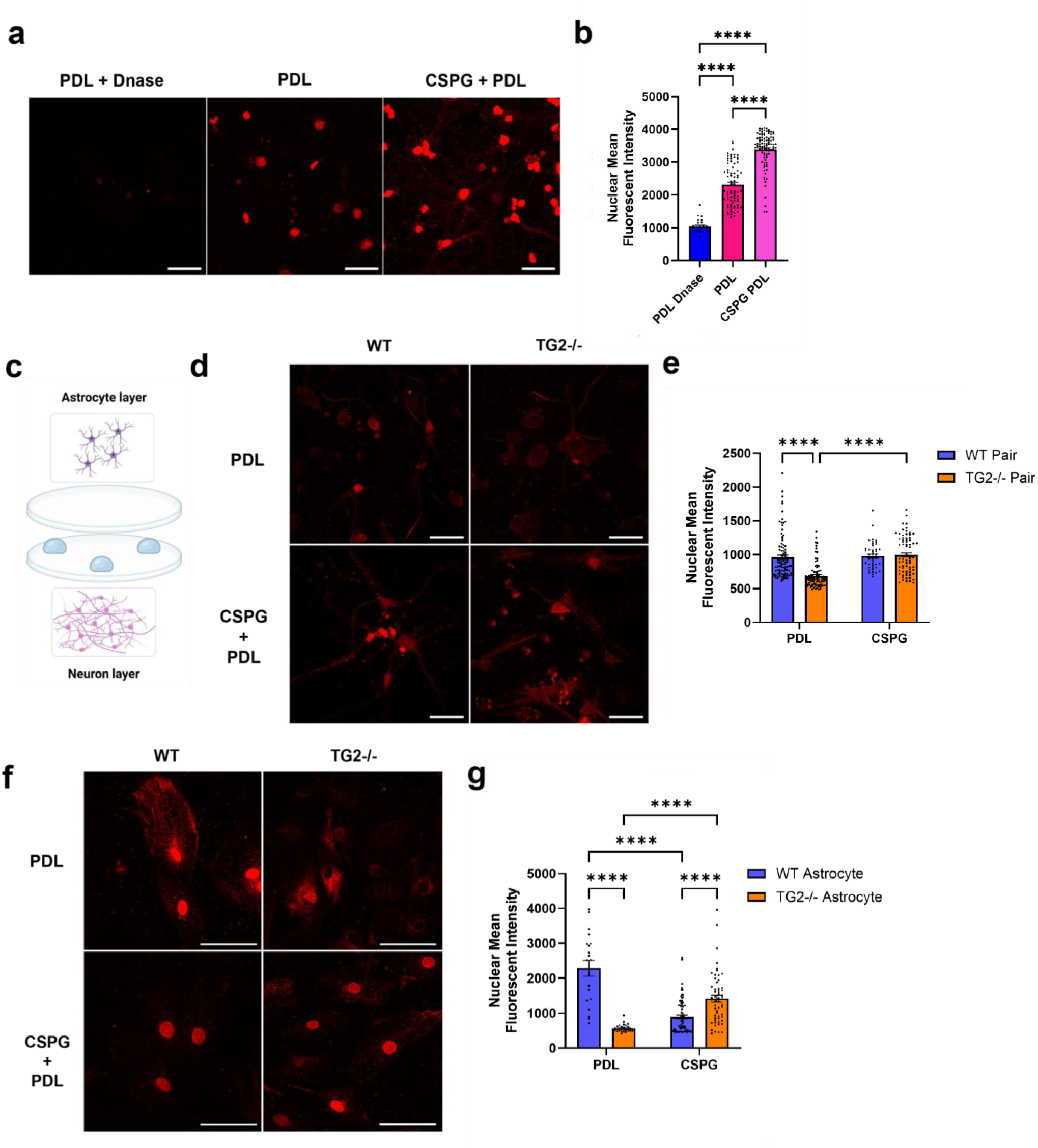
TG2-/- astrocytes better support redox balance in co-cultured neurons. Neuron (DIV 14) and astrocyte cultures (DIV 3-4) were immunolabeled for 8-oxo-guanine, a marker of DNA oxidation. **a)** Representative images of 8-oxo-guanine staining of primary neurons cultured on PDL, PDL + CSPGs (a growth-inhibitory substrate), or DNase negative control, and **b)** quantified nuclear mean intensity of 8-oxo-guanine immunofluorescence (n= 27-95, with 2 biological replicates, one-way ANOVA with Tukey’s multiple comparison test). **c)** Neuron-astrocyte co-culture design using two glass coverslips separated by paraffin pedestals to allow imaging of both compartments. **d)** Representative images of neurons paired with WT or TG2-/- astrocytes, and **e)** quantified 8-oxo-guanine signal (n= 47-98, with 3 biological replicates, two-way ANOVA with Šidák’s multiple comparison test). **f-g)** Representative images and quantifications of 8-oxo-guanine signal in WT or TG2-/- astrocytes that were co-cultured with neurons (n= 20-72, with 3 biological replicates, Šidák’s multiple comparison test). ****P < 0.0001. Data in bar graphs represent mean ± s.e.m. Scale bars in **a),d),** and **f)** = 40 μm.

**Extended Data Figure 7.**
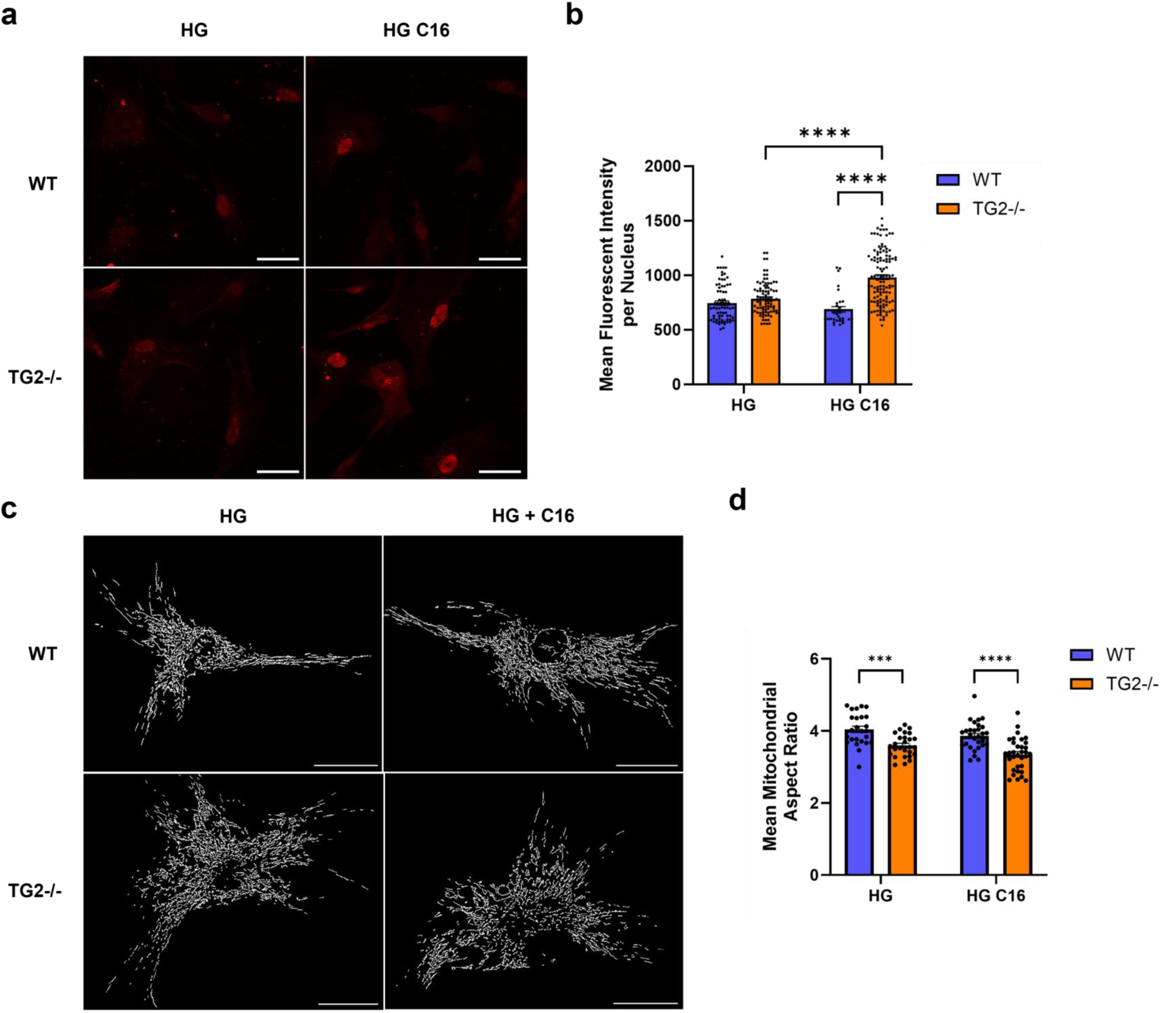
Oxidation status and mitochondrial morphology indicate that TG2-/- astrocytes upregulate lipid utilization. WT and TG2- /- primary astrocytes (DIV 3-4) were placed in high glucose media (HG, 33 mM glucose with 1 mM sodium pyruvate) or substrate-limited media (SL, 5.5 mM glucose without sodium pyruvate) with or without 50 mM palmitate (C16) for 24 hours. Oxidation status (8-oxo-guanine) and mitochondrial morphology (Tom20) were assessed as indicators of lipid utilization. **a)** Representative images of 8-oxo-guanine staining of astrocyte cultures across treatment groups (scale bars = 40 μm) and **b)** quantified nuclear mean intensity of 8- oxo-guanine immunofluorescence (n= 62- 215, with 2-3 biological replicates). **c)** Representative images of thresholded and binarized mitochondria derived from Tom20 staining across treatment groups (scale bars = 25 μm), and **d)** quantified mean mitochondrial aspect ratio for each group (n= 25-33, with 2 biological replicates). Data in bar graphs represent mean ± s.e.m. (two-way ANOVA with Šidák’s multiple comparison test; *P < 0.05, **P < 0.01, ***P < 0.001, ****P < 0.0001).

**Extended Data Figure 8.**
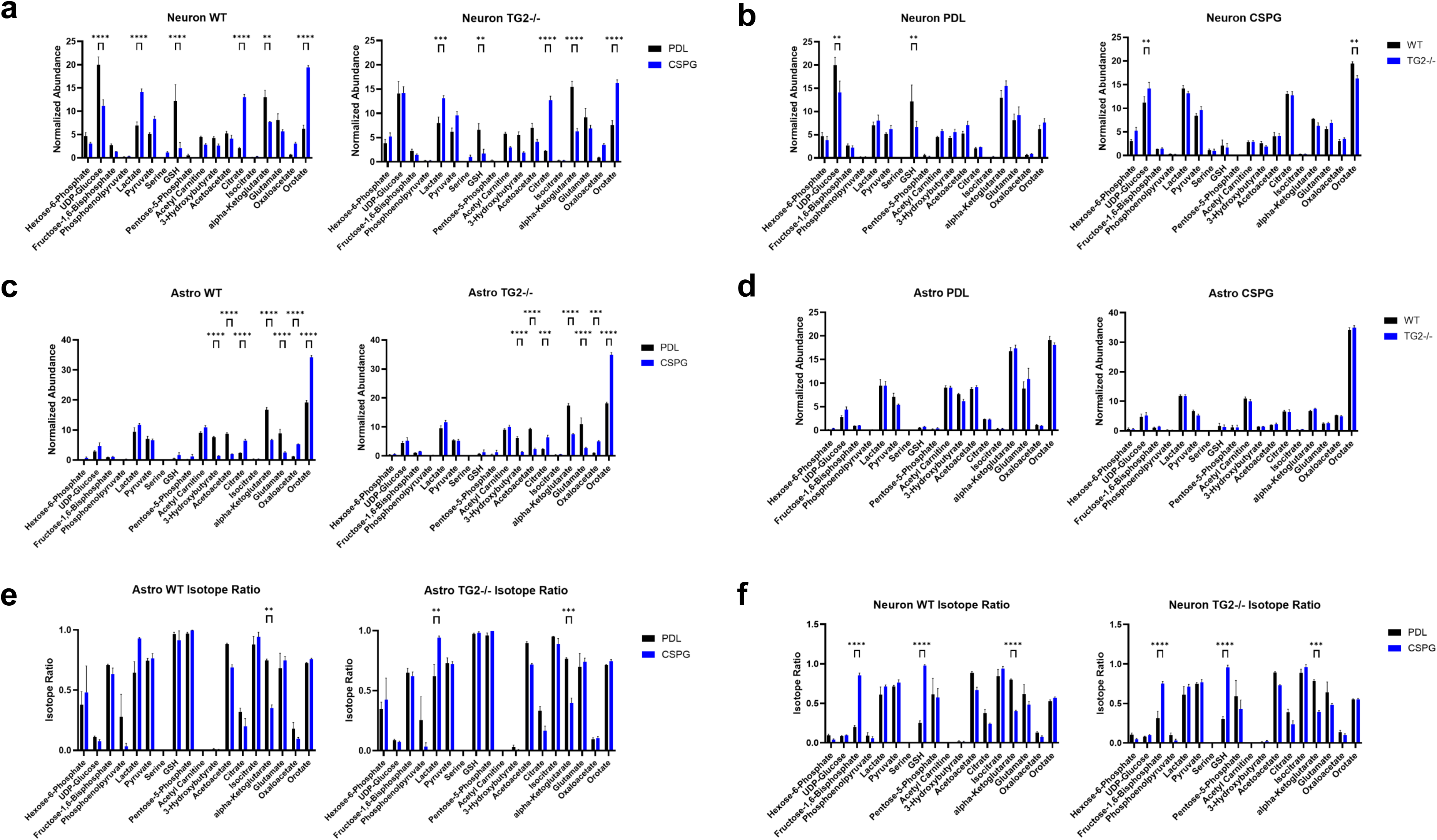
Metabolomics of astrocyte-neuron co-cultures following astrocyte incubation with lipid isotope. WT and TG2-/- primary astrocytes (DIV 3-4) were incubated with 50 mM uniformly labeled 13C palmitate for 24 hours. They were then paired with neurons (DIV 14) grown on CSPG+PDL or PDL matrices for 4 hours in artificial cerebral spinal fluid. **a-d)** LC-MS metabolomic analysis of neuron and astrocyte layers showing total abundance (abundance of labeled and unlabeled metabolites) across metabolites that were preselected for analysis. Results are separated for **(a,c)** matrix and **(b,d)** genotype comparisons. **e-f)** Isotope ratios (labeled metabolites/total abundance) of measured metabolites in astrocyte and neuron layers. Data in bar graphs represent mean ± s.e.m. (n= 4-5, with 2-3 biological replicates, two-way ANOVA with Šidák’s multiple comparison test; **P < 0.01, ***P < 0.001, ****P < 0.0001)

**Extended Data Figure 9.**
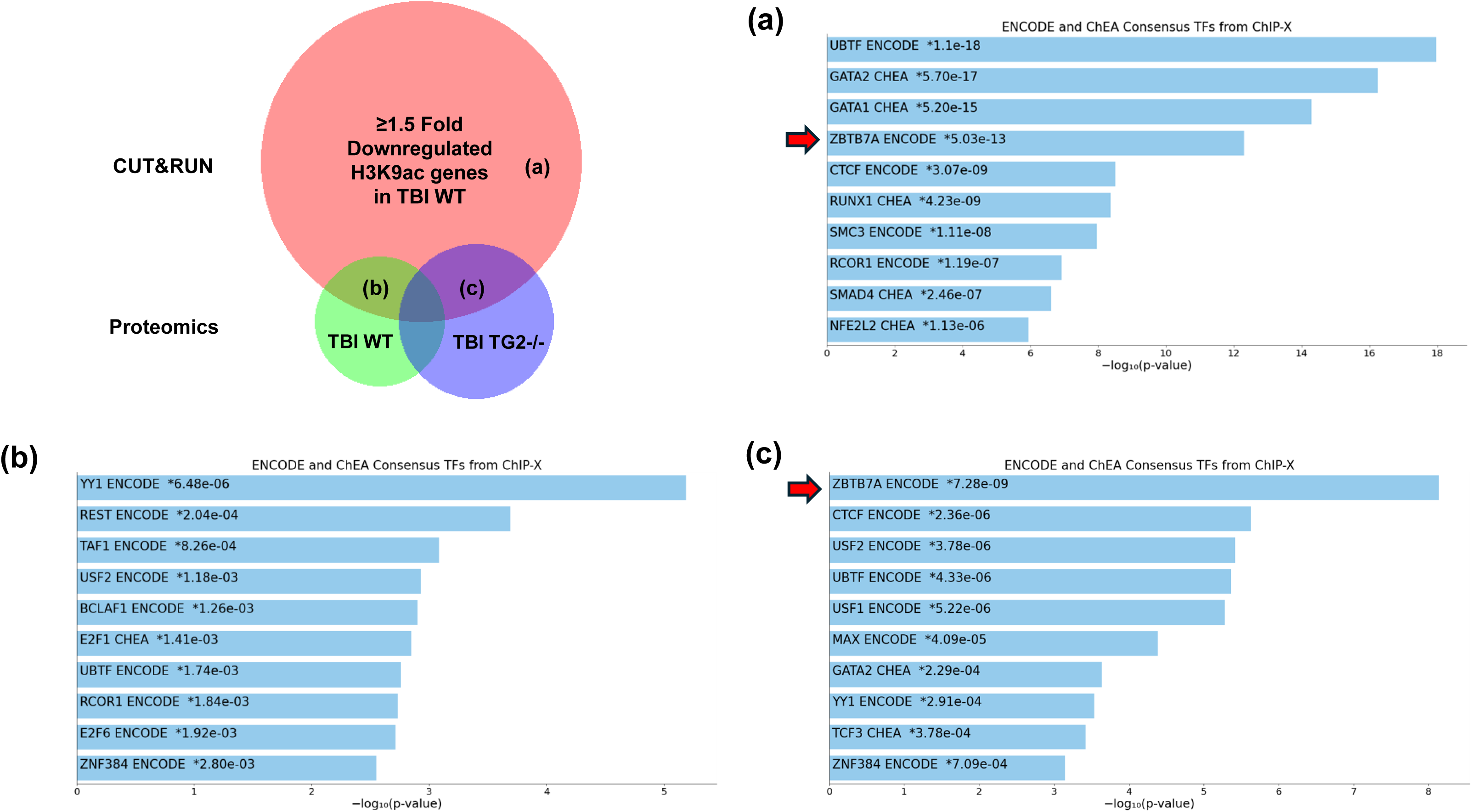
Transcription factor enrichment for genes with reduced H3K9ac signal after rmTBI identifies Zbtb7a among top categories. Venn diagram shows genes with ≥1.5-fold-change reduction in H3K9ac in TBI WT group that overlapped with differentially regulated proteins in TBI WT and TBI TG2-/-. Enrichr analysis using the ENCODE and ChEA databases was carried out on **a)** overall downregulated H3K9ac genes (5607 genes), and downregulated H3K9ac genes that overlapped with up- and down-regulated proteins in the **b)** TBI WT (381 genes) and **c)** TBI TG2-/- (496 genes) groups. Categories are ranked by p value. Asterisks represent categories with Benjamini Hochberg corrected p value <0.05. Venn diagram generated using BioVenn.

## References

1. Wu, L., et al., Global, regional, and national burdens of mild traumatic brain injuries from 1990 to 2019: findings from the Global Burden of Disease Study 2019 - a cross-sectional study. Int J Surg, 2025. 111(1): p. 160–170.

2. Dewan, M.C., et al., Estimating the global incidence of traumatic brain injury. J Neurosurg, 2019. 130(4): p. 1080–1097.

3. Rubiano, A.M., et al., Global neurotrauma research challenges and opportunities. Nature, 2015. 527(7578): p. S193–7.

4. Dery, J., et al., Prognostic factors for persistent symptoms in adults with mild traumatic brain injury: an overview of systematic reviews. Syst Rev, 2023. 12(1): p. 127.

5. Manley, G.T., et al., A new characterisation of acute traumatic brain injury: the NIH-NINDS TBI Classification and Nomenclature Initiative. Lancet Neurol, 2025. 24(6): p. 512–523.

6. Danna-Dos-Santos, A., et al., Author Correction: Long-term effects of mild traumatic brain injuries to oculomotor tracking performances and reaction times to simple environmental stimuli. Sci Rep, 2018. 8(1): p. 8174.

7. Madhok, D.Y., et al., Outcomes in Patients With Mild Traumatic Brain Injury Without Acute Intracranial Traumatic Injury. JAMA Netw Open, 2022. 5(8): p. e2223245.

8. Schwab, K., et al., Epidemiology and prognosis of mild traumatic brain injury in returning soldiers: A cohort study. Neurology, 2017. 88(16): p. 1571–1579.

9. Graham, A., et al., Mild Traumatic Brain Injuries and Future Risk of Developing Alzheimer’s Disease: Systematic Review and Meta-Analysis. J Alzheimers Dis, 2022. 87(3): p. 969–979.

10. Barnes, D.E., et al., Association of Mild Traumatic Brain Injury With and Without Loss of Consciousness With Dementia in US Military Veterans. JAMA Neurol, 2018. 75(9): p. 1055–1061.

11. Hiskens, M.I., et al., Repetitive mild traumatic brain injury-induced neurodegeneration and inflammation is attenuated by acetyl-L-carnitine in a preclinical model. Front Pharmacol, 2023. 14: p. 1254382.

12. Lehman, E.J., et al., Neurodegenerative causes of death among retired National Football League players. Neurology, 2012. 79(19): p. 1970–4.

13. Xu, X., et al., Repetitive mild traumatic brain injury in mice triggers a slowly developing cascade of long-term and persistent behavioral deficits and pathological changes. Acta Neuropathol Commun, 2021. 9(1): p. 60.

14. Lyons, D.N., et al., A Mild Traumatic Brain Injury in Mice Produces Lasting Deficits in Brain Metabolism. J Neurotrauma, 2018. 35(20): p. 2435–2447.

15. Barres, B.A., The mystery and magic of glia: a perspective on their roles in health and disease. Neuron, 2008. 60(3): p. 430–40.

16. Sofroniew, M.V. and H.V. Vinters, Astrocytes: biology and pathology. Acta Neuropathol, 2010. 119(1): p. 7–35.

17. Bonvento, G. and J.P. Bolanos, Astrocyte-neuron metabolic cooperation shapes brain activity. Cell Metab, 2021. 33(8): p. 1546–1564.

18. Bolanos, J.P. and P.J. Magistretti, The neuron-astrocyte metabolic unit as a cornerstone of brain energy metabolism in health and disease. Nat Metab, 2025. 7(12): p. 2414–2423.

19. Escartin, C., et al., Reactive astrocyte nomenclature, definitions, and future directions. Nat Neurosci, 2021. 24(3): p. 312–325.

20. Myer, D.J., et al., Essential protective roles of reactive astrocytes in traumatic brain injury. Brain, 2006. 129(Pt 10): p. 2761–72.

21. Liddelow, S.A., et al., Neurotoxic reactive astrocytes are induced by activated microglia. Nature, 2017. 541(7638): p. 481–487.

22. He, L., et al., The role of astrocyte in neuroinflammation in traumatic brain injury. Biochim Biophys Acta Mol Basis Dis, 2024. 1870(3): p. 166992.

23. Jamjoom, A.A.B., et al., The synapse in traumatic brain injury. Brain, 2021. 144(1): p. 18–31.

24. Xiong, X.Y., Y. Tang, and Q.W. Yang, Metabolic changes favor the activity and heterogeneity of reactive astrocytes. Trends Endocrinol Metab, 2022. 33(6): p. 390–400.

25. Tatsukawa, H. and K. Hitomi, Role of Transglutaminase 2 in Cell Death, Survival, and Fibrosis. Cells, 2021. 10(7).

26. Colak, G. and G.V. Johnson, Complete transglutaminase 2 ablation results in reduced stroke volumes and astrocytes that exhibit increased survival in response to ischemia. Neurobiol Dis, 2012. 45(3): p. 1042–50.

27. Quinn, B.R., L. Yunes-Medina, and G.V.W. Johnson, Transglutaminase 2: Friend or foe? The discordant role in neurons and astrocytes. J Neurosci Res, 2018. 96(7): p. 1150–1158.

28. Monteagudo, A., et al., Depletion of astrocytic transglutaminase 2 improves injury outcomes. Mol Cell Neurosci, 2018. 92: p. 128–136.

29. Folk, J.E. and J.S. Finlayson, The epsilon-(gamma-glutamyl)lysine crosslink and the catalytic role of transglutaminases. Adv Protein Chem, 1977. 31: p. 1–133.

30. Folk, J.E., Transglutaminases. Annu Rev Biochem, 1980. 49: p. 517–31.

31. Akimov, S.S., et al., Tissue transglutaminase is an integrin-binding adhesion coreceptor for fibronectin. J Cell Biol, 2000. 148(4): p. 825–38.

32. Kanchan, K., M. Fuxreiter, and L. Fesus, Physiological, pathological, and structural implications of non-enzymatic protein-protein interactions of the multifunctional human transglutaminase 2. Cell Mol Life Sci, 2015. 72(16): p. 3009–35.

33. Lesort, M., et al., Distinct nuclear localization and activity of tissue transglutaminase. J Biol Chem, 1998. 273(20): p. 11991–4.

34. Filiano, A.J., et al., Transglutaminase 2 protects against ischemic stroke. Neurobiol Dis, 2010. 39(3): p. 334–43.

35. Yunes-Medina, L., J. Feola, and G.V.W. Johnson, Subcellular localization patterns of transglutaminase 2 in astrocytes and neurons are differentially altered by hypoxia. Neuroreport, 2017. 28(18): p. 1208–1214.

36. Piacentini, M., et al., Characterization of distinct sub-cellular location of transglutaminase type II: changes in intracellular distribution in physiological and pathological states. Cell Tissue Res, 2014. 358(3): p. 793–805.

37. Emerson, J., et al., Deletion of Transglutaminase 2 from Mouse Astrocytes Significantly Improves Their Ability to Promote Neurite Outgrowth on an Inhibitory Matrix. Int J Mol Sci, 2023. 24(7).

38. McConoughey, S.J., et al., Inhibition of transglutaminase 2 mitigates transcriptional dysregulation in models of Huntington disease. EMBO Mol Med, 2010. 2(9): p. 349–70.

39. Farrelly, L.A., et al., Histone serotonylation is a permissive modification that enhances TFIID binding to H3K4me3. Nature, 2019. 567(7749): p. 535–539.

40. Zheng, Q., et al., Bidirectional histone monoaminylation dynamics regulate neural rhythmicity. Nature, 2025. 637(8047): p. 974–982.

41. Kuo, T.F., H. Tatsukawa, and S. Kojima, New insights into the functions and localization of nuclear transglutaminase 2. FEBS J, 2011. 278(24): p. 4756–67.

42. Delgado, T., et al., Pharmacological Inhibition of Astrocytic Transglutaminase 2 Facilitates the Expression of a Neurosupportive Astrocyte Reactive Phenotype in Association with Increased Histone Acetylation. Biomolecules, 2024. 14(12).

43. Delgado T, J.G., Transglutaminase 2 in Neurological Conditions, in Transglutaminase Fundamentals and Applications, Y. In Zhang, & Simpson, BK, Editor. 2024, Elsevier Publishing. p. 107–122.

44. Elahi, A., et al., Deletion or Inhibition of Astrocytic Transglutaminase 2 Promotes Functional Recovery after Spinal Cord Injury. Cells, 2021. 10(11).

45. Oft, H.C., D.W. Simon, and D. Sun, New insights into metabolism dysregulation after TBI. J Neuroinflammation, 2024. 21(1): p. 184.

46. Olatona, O.A., et al., Mitochondria: the hidden engines of traumatic brain injury-driven neurodegeneration. Front Cell Neurosci, 2025. 19: p. 1570596.

47. Crum, A.B., et al., Development of a Modified Weight-Drop Apparatus for Closed-Skull, Repetitive Mild Traumatic Brain Injuries in a Mouse Model. eNeuro, 2026. 13(1).

48. Hamm, R.J., Neurobehavioral assessment of outcome following traumatic brain injury in rats: an evaluation of selected measures. J Neurotrauma, 2001. 18(11): p. 1207–16.

49. Charmant, J., Kinovea (version 2025.1). [Online], (Available): p. https://kinovea.readthedocs.io/en/latest/index.html.

50. Zang, Y.F., et al., Altered baseline brain activity in children with ADHD revealed by resting-state functional MRI. Brain Dev, 2007. 29(2): p. 83–91.

51. Arefin, T.M., et al., Towards reliable reconstruction of the mouse brain corticothalamic connectivity using diffusion MRI. Neuroimage, 2023. 273: p. 120111.

52. Zuidema, T.R., et al., Longitudinal Associations of Clinical and Biochemical Head Injury Biomarkers With Head Impact Exposure in Adolescent Football Players. JAMA Netw Open, 2023. 6(5): p. e2316601.

53. Bazarian, J.J., et al., A Pilot Study Investigating the Use of Serum Glial Fibrillary Acidic Protein to Monitor Changes in Brain White Matter Integrity After Repetitive Head Hits During a Single Collegiate Football Game. J Neurotrauma, 2024. 41(13-14): p. 1597–1608.

54. Abou-El-Hassan, H., et al., Astrocyte activation persists one year after TBI: a dynamic shift from inflammation to neurodegeneration. Commun Biol, 2025. 8(1): p. 1745.

55. Munoz-Ballester, C., et al., Mild Traumatic Brain Injury-Induced Disruption of the Blood-Brain Barrier Triggers an Atypical Neuronal Response. Front Cell Neurosci, 2022. 16: p. 821885.

56. Tiana, M., et al., The SIN3A histone deacetylase complex is required for a complete transcriptional response to hypoxia. Nucleic Acids Res, 2018. 46(1): p. 120–133.

57. Icardi, L., et al., The Sin3a repressor complex is a master regulator of STAT transcriptional activity. Proc Natl Acad Sci U S A, 2012. 109(30): p. 12058–63.

58. Gates, L.A., et al., Acetylation on histone H3 lysine 9 mediates a switch from transcription initiation to elongation. J Biol Chem, 2017. 292(35): p. 14456– 14472.

59. Nessel, I. and A.T. Michael-Titus, Lipid profiling of brain tissue and blood after traumatic brain injury: A review of human and experimental studies. Semin Cell Dev Biol, 2021. 112: p. 145–156.

60. Waagepetersen, H.S., et al., Demonstration of pyruvate recycling in primary cultures of neocortical astrocytes but not in neurons. Neurochem Res, 2002. 27(11): p. 1431–7.

61. Bak, L.K., et al., Complex glutamate labeling from [U-13C]glucose or [U- 13C]lactate in co-cultures of cerebellar neurons and astrocytes. Neurochem Res, 2007. 32(4-5): p. 671–80.

62. Sakamoto, K., et al., Glycan sulfation patterns define autophagy flux at axon tip via PTPRsigma-cortactin axis. Nat Chem Biol, 2019. 15(7): p. 699–709.

63. Sakamoto, K., T. Ozaki, and K. Kadomatsu, Axonal Regeneration by Glycosaminoglycan. Front Cell Dev Biol, 2021. 9: p. 702179.

64. Ioannou, M.S., et al., Neuron-Astrocyte Metabolic Coupling Protects against Activity-Induced Fatty Acid Toxicity. Cell, 2019. 177(6): p. 1522–1535 e14.

65. Ngo, J., et al., Mitochondrial morphology controls fatty acid utilization by changing CPT1 sensitivity to malonyl-CoA. EMBO J, 2023. 42(11): p. e111901.

66. Morant-Ferrando, B., et al., Fatty acid oxidation organizes mitochondrial supercomplexes to sustain astrocytic ROS and cognition. Nat Metab, 2023. 5(8): p. 1290–1302.

67. Liesa, M. and O.S. Shirihai, Mitochondrial dynamics in the regulation of nutrient utilization and energy expenditure. Cell Metab, 2013. 17(4): p. 491–506.

68. Mao, C., et al., DHODH-mediated ferroptosis defence is a targetable vulnerability in cancer. Nature, 2021. 593(7860): p. 586–590.

69. Padalko, V., F. Posnik, and M. Adamczyk, Mitochondrial Aconitase and Its Contribution to the Pathogenesis of Neurodegenerative Diseases. Int J Mol Sci, 2024. 25(18).

70. Yan, L.J., R.L. Levine, and R.S. Sohal, Oxidative damage during aging targets mitochondrial aconitase. Proc Natl Acad Sci U S A, 1997. 94(21): p. 11168–72.

71. Bazarian, J.J., et al., Persistent, long-term cerebral white matter changes after sports-related repetitive head impacts. PLoS One, 2014. 9(4): p. e94734.

72. Hayes, J.P., E.D. Bigler, and M. Verfaellie, Traumatic Brain Injury as a Disorder of Brain Connectivity. J Int Neuropsychol Soc, 2016. 22(2): p. 120–37.

73. Zhou, Y., et al., Default-mode network disruption in mild traumatic brain injury. Radiology, 2012. 265(3): p. 882–92.

74. Lane, A.N. and T.W. Fan, Regulation of mammalian nucleotide metabolism and biosynthesis. Nucleic Acids Res, 2015. 43(4): p. 2466–85.

75. Jackson, T.C. and P.M. Kochanek, RNA Binding Motif 5 (RBM5) in the CNS- Moving Beyond Cancer to Harness RNA Splicing to Mitigate the Consequences of Brain Injury. Front Mol Neurosci, 2020. 13: p. 126.

76. Sardar, D., et al., Induction of astrocytic Slc22a3 regulates sensory processing through histone serotonylation. Science, 2023. 380(6650): p. eade0027.

77. Gupta, S., et al., Emerging role of ZBTB7A as an oncogenic driver and transcriptional repressor. Cancer Lett, 2020. 483: p. 22–34.

78. Ramos Pittol, J.M., et al., Zbtb7a is a transducer for the control of promoter accessibility by NF-kappa B and multiple other transcription factors. PLoS Biol, 2018. 16(5): p. e2004526.

79. Choi, W.I., et al., Proto-oncogene FBI-1 represses transcription of p21CIP1 by inhibition of transcription activation by p53 and Sp1. J Biol Chem, 2009. 284(19): p. 12633–44.

80. Rodriguez-Campuzano, A.G., et al., Yin Yang 1: Function, Mechanisms, and Glia. Neurochem Res, 2025. 50(2): p. 96.

81. Schindelin, J., et al., Fiji: an open-source platform for biological-image analysis. Nat Methods, 2012. 9(7): p. 676–82.

82. Ji, C., et al., Nuclear transglutaminase 2 directly regulates expression of cathepsin S in rat cortical neurons. Eur J Neurosci, 2018. 48(9): p. 3043–3051.

83. Hainer, S.J., et al., Profiling of Pluripotency Factors in Single Cells and Early Embryos. Cell, 2019. 177(5): p. 1319–1329 e11.

84. Skene, P.J. and S. Henikoff, An efficient targeted nuclease strategy for high-resolution mapping of DNA binding sites. Elife, 2017. 6.

85. Yeh, S.Y., et al., Cell Type-Specific Whole-Genome Landscape of DeltaFOSB Binding in the Nucleus Accumbens After Chronic Cocaine Exposure. Biol Psychiatry, 2023. 94(5): p. 367–377.

86. Langmead, B. and S.L. Salzberg, Fast gapped-read alignment with Bowtie 2. Nat Methods, 2012. 9(4): p. 357–9.

87. Li, H., et al., The Sequence Alignment/Map format and SAMtools. Bioinformatics, 2009. 25(16): p. 2078–9.

88. Ramirez, F., et al., deepTools: a flexible platform for exploring deep-sequencing data. Nucleic Acids Res, 2014. 42(Web Server issue): p. W187–91.

89. Huang da, W., B.T. Sherman, and R.A. Lempicki, Systematic and integrative analysis of large gene lists using DAVID bioinformatics resources. Nat Protoc, 2009. 4(1): p. 44–57.

90. Chen, E.Y., et al., Enrichr: interactive and collaborative HTML5 gene list enrichment analysis tool. BMC Bioinformatics, 2013. 14: p. 128.

91. Kuleshov, M.V., et al., Enrichr: a comprehensive gene set enrichment analysis web server 2016 update. Nucleic Acids Res, 2016. 44(W1): p. W90–7.

92. Xie, Z., et al., Gene Set Knowledge Discovery with Enrichr. Curr Protoc, 2021. 1(3): p. e90.

93. Milliken, A.S., et al., Distinct effects of intracellular vs. extracellular acidic pH on the cardiac metabolome during ischemia and reperfusion. J Mol Cell Cardiol, 2023. 174: p. 101–114.

94. Aittokallio, T., Dealing with missing values in large-scale studies: microarray data imputation and beyond. Brief Bioinform, 2010. 11(2): p. 253–64.

95. Arefin, T.M., et al., Remodeling of Sensorimotor Brain Connectivity in Gpr88- Deficient Mice. Brain Connect, 2017. 7(8): p. 526–540.

96. Mechling, A.E., et al., Deletion of the mu opioid receptor gene in mice reshapes the reward-aversion connectome. Proc Natl Acad Sci U S A, 2016. 113(41): p. 11603–11608.

97. Hennig, J., A. Nauerth, and H. Friedburg, RARE imaging: a fast imaging method for clinical MR. Magn Reson Med, 1986. 3(6): p. 823–33.

98. Liu, Y., et al., An open database of resting-state fMRI in awake rats. Neuroimage, 2020. 220: p. 117094.

99. Arefin, T.M., et al., Dissecting the Strain and Sex Specific Connectome Signatures of Unanesthetized C57BL/6J and DBA/2J Mice Using Magnetic Resonance Imaging. Genes Brain Behav, 2026. 25(4): p. e70060.

100. Power, J.D., et al., Spurious but systematic correlations in functional connectivity MRI networks arise from subject motion. Neuroimage, 2012. 59(3): p. 2142–54.

101. Arefin, T.M., et al., Macroscopic Structural and Connectome Mapping of the Mouse Brain Using Diffusion Magnetic Resonance Imaging. Bio Protoc, 2021. 11(22): p. e4221.

102. Ben Youss, Z., et al., Open-source versatile 3D-print animal conditioning platform design for in vivo preclinical brain imaging in awake mice and anesthetized mice and rats. Lab Anim (NY), 2024. 53(2): p. 33–42.

103. Jiang, H., et al., DtiStudio: resource program for diffusion tensor computation and fiber bundle tracking. Comput Methods Programs Biomed, 2006. 81(2): p. 106– 16.

104. Arefin, T.M., et al., Sex-specific signatures of GLP-1 and amylin on resting state brain activity and functional connectivity in awake rats. Neuropharmacology, 2025. 269: p. 110348.

105. Mori, S. and J. Zhang, Principles of diffusion tensor imaging and its applications to basic neuroscience research. Neuron, 2006. 51(5): p. 527–39.

106. Basser, P.J. and C. Pierpaoli, Microstructural and physiological features of tissues elucidated by quantitative-diffusion-tensor MRI. J Magn Reson B, 1996. 111(3): p. 209–19.

